# Pleotropic effects of PPARD accelerate colorectal tumor progression and invasion

**DOI:** 10.1101/325001

**Authors:** Yi Liu, Yasunori Deguchi, Rui Tian, Daoyan Wei, Weidong Chen, Min Xu, Ling Wu, Fuyao Liu, Shen Gao, Jonathan C. Jaoude, Sarah P. Chrieki, Micheline J. Moussalli, Mihai Gagea, Jeffrey S. Morris, Russell Broaddus, Xiangsheng Zuo, Imad Shureiqi

**Affiliations:** Department of Gastrointestinal Medical Oncology, The University of Texas MD Anderson Cancer Center, Houston, Texas 77030, USA; Department of Gastroenterology, Hepatology, and Nutrition,The University of Texas MD Anderson Cancer Center, Houston, Texas 77030, USA; Department of Pathology, The University of Texas MD Anderson Cancer Center, Houston, Texas 77030, USA; Department of Veterinary Medicine & Surgery, The University of Texas MD Anderson Cancer Center, Houston, Texas 77030, USA; Department of Biostatistics, The University of Texas MD Anderson Cancer Center, Houston, Texas 77030, USA; Department of Biliary-Pancreatic Surgery, Affiliated Tongji Hospital, Tongji Medical College, Huazhong University of Science and Technology, Wuhan, Hubei, 430030, China

## Abstract

Colorectal carcinogenesis (CRC) progression requires additional molecular mechanisms to APC mutations/aberrant β-catenin signaling. PPARD is a druggable ligand-activated nuclear receptor that regulates essential genes involved in cell fate. PPARD is upregulated in intestinal epithelial cells (IECs) of human colorectal adenomas and adenocarcinomas. The mechanistic significance of PPARD upregulation in CRC remains unknown. Here we show that targeted PPARD overexpression in IECs of mice strongly augmented β-catenin activation via BMP7/TAK1 signaling, promoted intestinal tumorigenesis in Apc^min^ mice, and accelerated CRC progression and invasiveness in mice with IEC-targeted Apc^Δ580^ mutation. Human CRC invasive fronts had higher PPARD expression than their paired adenomas. A PPARD agonist (GW501516) enhanced APC^Δ580^ mutation-driven CRC, while a PPARD antagonist (GSK3787) suppressed it. Functional proteomics analyses and subsequent validation studies uncovered PPARD upregulation of multiple pro-invasive pathways that drive CRC progression (e.g. PDGFRβ, AKT1, CDK1 and EIF4G1). Our results identify novel mechanisms by which PPARD promotes CRC invasiveness and provide the rational for the development of PPARD antagonists to suppress CRC.

## Introduction

APC mutations activate aberrant β-catenin signaling to drive colorectal tumorigenesis(1). However, colorectal cancer (CRC) progression, especially invasiveness, requires additional molecular mechanisms (2, 3). The presence of CRC invasion in colorectal polyps profoundly worsen patients’ outcomes (4). Identification of critical invasiveness-regulating mechanisms can therefore open opportunities for molecular targeting.

The ligand-activated nuclear receptor peroxisome proliferator-activated receptor-δ/β (PPARD) has pleiotropic effects on cell homeostasis (5). PPARD is upregulated in human colorectal polyps and CRC (6-9) and has been identified as a downstream target of the β-catenin/TCF4 complex in CRC cell lines (6). However, this proposed mechanistic link between β-catenin and PPARD has been questioned on the basis of in vitro and in vivo data(10). More importantly, germline PPARD knockout (KO) in Apc^min^ mice produced conflicting results, both increasing (11) and decreasing (12)intestinal tumorigenesis. Nevertheless, other studies suggested that PPARD increases β-catenin activation in human cholangiocarcinoma cells (13) and in normal osteocytes(14). Recently, a high-fat diet was reported to increase β-catenin activation via PPARD in progenitor intestinal cells of Apc^min^ mice (15). Clearly, the role of PPARD in colorectal tumorigenesis, especially in relation to APC and aberrant β-catenin activation, remains highly controversial (16, 17). Filling this knowledge gap is important because PPARD is a druggable protein for which agonists and antagonists have been developed. Although the clinical testing and pharmaceutical development of PPARD agonists by large pharmaceutical companies to treat noncancerous conditions (e.g., obesity) has been halted in many instances, these agents (e.g., cardarine [GW501516]) are still sold on the internet black market to athletes wishing to enhance muscle endurance. Therefore, preclinical data clarifying the role of PPARD in CRC are urgently needed to educate the public about the potential risk of promoting colorectal tumorigenesis with PPARD agonists.

We therefore tested PPARD’s effects on aberrant β-catenin activation-driven colon tumorigenesis using murine genetic models of human CRC with representative APC mutations(18) with concomitant PPARD overexpression in intestinal epithelial cells (IECs). Our data showed that PPARD strongly enhanced aberrant β-catenin activation and more importantly it robustly activated CRC proinvasive pathways to promote CRC progression and invasion.

## Results

### PPARD activates β-catenin in IECs

Aberrant β-catenin activation via APC mutations upregulates PPARD (6). We therefore tested whether PPARD upregulation is sufficient to increase β-catenin activation independently of APC mutations by using PD mice with wild-type (WT) APC (Supplementary Fig. 1a). PPARD overexpression markedly increased active β-catenin protein levels in IECs (Fig.1a). The PPARD agonist GW501516 augmented the effects of PPARD in PD mice compared to their WT littermates (Supplementary Fig. 1b). PPARD overexpression also significantly increased mRNA levels of Axin2 and cyclinD1, downstream targets of active β-catenin, in IECs (Fig. 1b). Aberrant β-catenin activation expands the dedifferentiated intestinal stem cell population to promote colorectal tumorigenesis(19). Intestinal organoid formation, especially in its primitive spheroid form, was markedly higher when IECs were derived from PD mice than from their WT littermates (Fig. 1c).

**Fig 1.**
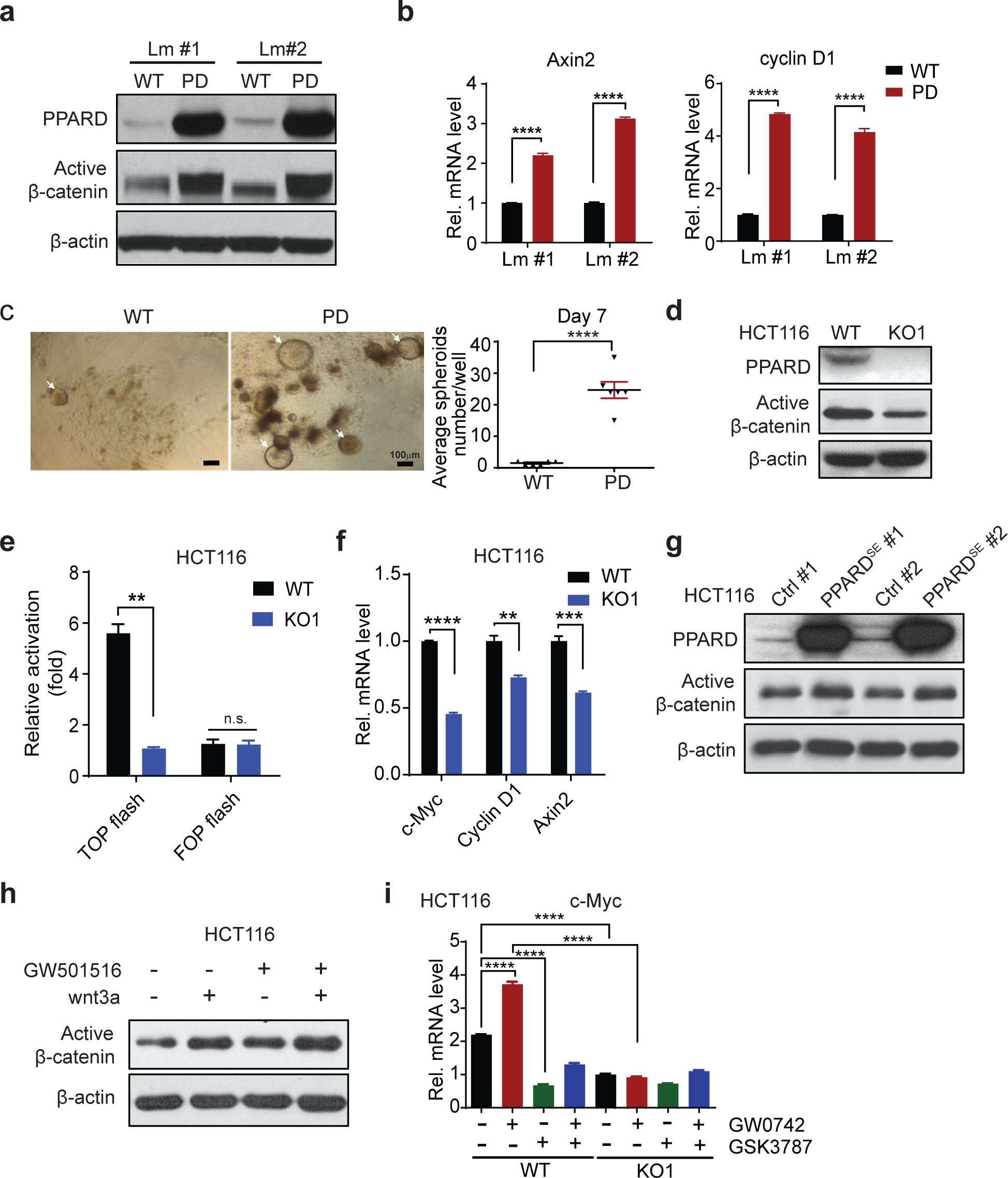
PPARD activates β-catenin signaling in intestinal epithelial cells. **(a, b)** Active β-catenin protein levels **(a)** and mRNA levels of β-catenin target-genes (Axin2 and cyclin D1) **(b)** in intestinal epithelial cells (IECs) of PD mice and WT littermates at age 10 weeks. Lm indicate littermate. **(c)** The organoid-initiating capacity of IECs derived from PD mice and WT littermates (n = 6 mice/group). Photographs of primary organoids (left); primary organoid counts (right). White arrows indicate individual organoids. **(d-f)** Active β-catenin protein levels **(d)**, β-catenin transcriptional activity (**e**, TOP/FOP Flash luciferase assay) and mRNA levels of Axin2, c-Myc and cyclin D1 **(f)** in HCT-116 cells without (WT) or with PPARD knockout (KO1). **(g, h)** Active β-catenin protein levels of stably transfected HCT-116 cells with control (Ctrl; # represents clone number) or PPARD overexpression plasmid (PPARD^SE^) **(g)** and HCT-116 cells treated with wnt3a (100 μM) and/or PPARD agonist GW501516 (1 μM) for 24 h **(h)**. **(I)** c-Myc mRNA expression levels in HCT-116 WT and KO1 cells treated with PPARD agonist GW0742 (1 μM) and/or PPARD antagonist GSK3787 (1 μM) for 24 h. The experiments were independently replicated three times. Error bars: mean ± s.e.m. \*\**P* < 0.01, \*\*\**P* < 0.001, \*\*\*\**P* < 0.0001.

We further examined whether PPARD activates β-catenin independently of APC mutations using HCT-116 human CRC cells, which have WT APC but a heterozygous β-catenin-activating mutation(1). Genetic deletion of PPARD in HCT-116 cells (KO1) significantly decreased active β-catenin protein expression (Fig. 1d), transcriptional activity (measured by a TCF/LEF luciferase reporter assay [TOP-Flash/FOP-Flash]) (Fig. 1e), and target gene expression (Axin2, cyclin D1, and c-Myc) (Fig. 1f). In contrast, PPARD overexpression in HCT-116 cells increased active β-catenin protein levels (Fig. 1g). Furthermore, the PPARD agonist GW501516 significantly increased active β-catenin protein levels and augmented Wnt3a-induced β-catenin activation in HCT-116 cells (Fig. 1h). Another PPARD agonist, GW0742, increased c-Myc mRNA expression in HCT-116 parental cells (WT) but not in KO1 cells (Fig. 1i), whereas the PPARD antagonist GSK3787 decreased c-Myc mRNA expression even in GW0742-treated cells (Fig. 1i). We confirmed PPARD activation by GW0742 and inhibition by GSK3787 in HCT-116 cells by measuring expression of angiopoietin-like 4 (AngPTL4) (Supplementary Fig. 1c), a well-established PPARD target gene.

### PPARD enhances aberrant β-catenin signaling in IECs with APC mutations

Apc^min^ mice mainly develop small intestinal adenomas, unlike humans, in whom intestinal tumorigenesis is essentially colorectal and can progress to invasive cancer. In contrast to Apc^min^ mice, targeting the APC mutation into the intestine via CDX2 promoter-driven Cre recombinase expression to produce a codon 580 frame-shift mutation, as in Apc^Δ580^-flox; CDX2-Cre (Apc^Δ580^) mice, results in the development of CRCs.(18) Breeding Apc^Δ580^ with PD mice generated Apc^Δ580^-PD mice (Supplementary Fig. 1d), which had significantly higher expression levels of PPARD (Supplementary Fig. 1e and Fig. 2a), active β-catenin protein (Fig. 2a,b and Supplementary Fig. 1f), and mRNA of β-catenin downstream targets Axin-2, c-Myc, and cyclin D1 in IECs than did Apc^Δ580^ mice (Fig. 2c). The PPARD agonist GW501516 also significantly increased active β-catenin protein expression in the IECs of Apc^Δ580^ mice (Fig. 2d, e). Apc^Δ580^-PD mice and GW501516-treated Apc^Δ580^ mice also had significantly longer colonic crypt proliferative zones than control Apc^Δ580^ mice (Fig. 2f,g).

**Fig 2.**
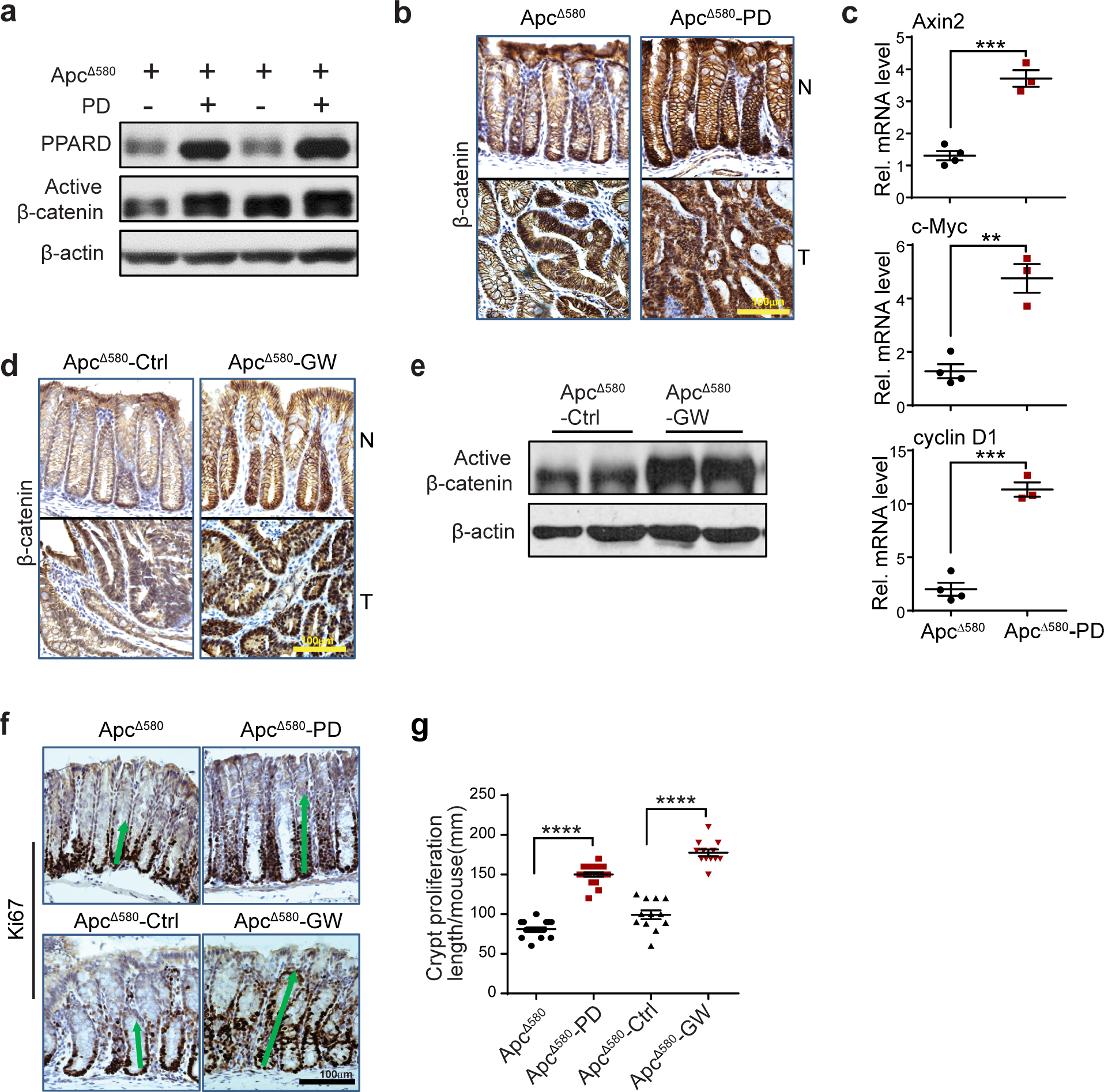
PPARD enhances intestinal β-catenin signaling in Apc^Δ580^ mutant mice. **(a-c)** Active β-catenin protein expression in intestinal epithelial cells (IECs) **(a)**; immunohistochemistry (IHC) localization of β-catenin in colonic normal and tumor tissues **(b)**; and Axin2, c-Myc and cyclin D1 mRNA expression levels in normal IECs **(c)** of Apc^Δ580^ and Apc^Δ580^-PD mice at age 14 weeks. **(d, e)** Apc^Δ580^ mice were fed a diet containing the PPARD agonist GW501516 (50 mg/kg) (GW) or a control diet (Ctrl) for 10 weeks, then evaluated for β-catenin expression by IHC **(d)** and active β-catenin protein expression **(e)** as described in panels a and b. **(f, g)** Representative images of Ki-67 IHC **(f)** and corresponding colonic crypt proliferation zone lengths **(g)** of normal colons of mice as described in panels a and d. Error bars: mean ± s.e.m. \*\**P <* 0.01, \*\*\**P <* 0.001, **** *P <* 0.0001.

### PPARD activates β-catenin via BMP7/TAK1 signaling in IECs

To determine the molecular mechanisms by which PPARD enhanced β-catenin activation, we used comparative transcriptome profile analyses (RNA sequencing) of HCT-116-WT and KO1 (HCT-116 PPARD KO) cells (9). These analyses revealed that bone morphogenetic protein 7 (BMP7) mRNA expression was 11-fold higher in WT cells than in KO1 cells (Supplementary Fig. 2a). Independent quantitative real-time polymerase chain reaction (qRT-PCR) measurements confirmed this finding (Fig. 3a). BMP7 phosphorylates mitogen-activated protein kinase kinase kinase 7 (TAK1) to activate β-catenin(20). BMP7 and phosphorylated TAK1 (p-TAK1) protein levels were lower in HCT-116 KO1 cells than in WT cells (Fig. 3b). PPARD overexpression by lentivirus transduction in SW480 colon cancer cells with low BMP7 expression significantly increased expression of BMP7 mRNA (Fig. 3c), BMP7 protein, p-TAK1, phosphorylated P38 (p-p38), and active β-catenin protein (Fig. 3d). We next examined whether PPARD transcriptionally regulates BMP7 expression via binding to a potential PPARD binding site (pPDBS) in the BMP7 promoter (Fig. 3e). An in silico search using Genomatix MatInspector online software identified a pPDBS in the BMP7 promoter region near the transcription start site (Fig. 3e). PPARD overexpression in SW480 cells significantly increased PPARD binding (measured by a chromatin immunoprecipitation/quantitative PCR assay) to this pPDBS in the BMP7 promoter (Fig. 3f).

**Fig 3.**
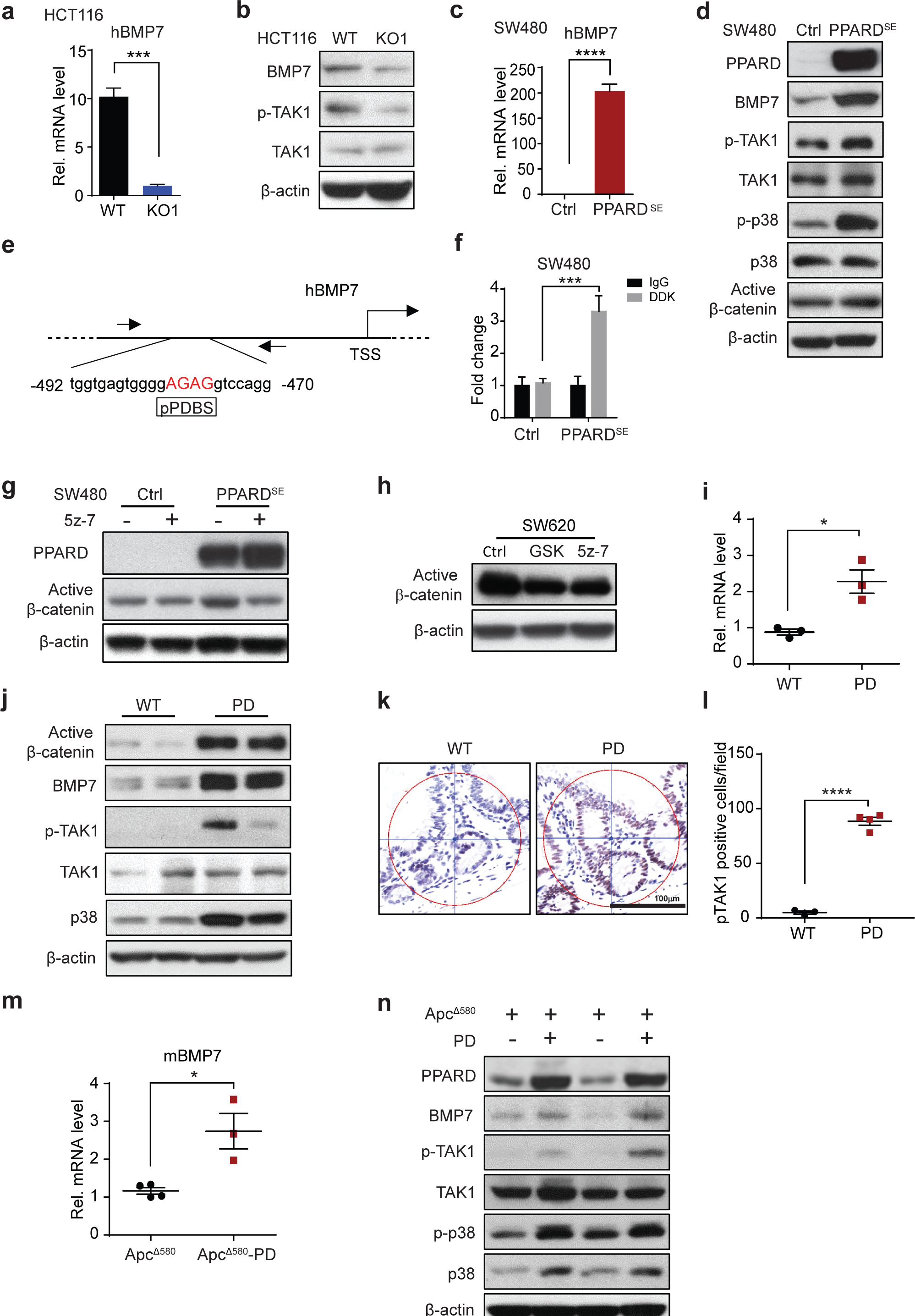
PPARD activates BMP7/TAK1/β-catenin signaling in intestinal epithelial cells. **(a)** BMP7 mRNA expression levels in WT or PPARD knockout (KO1) HCT-116 cells. **(b)** Protein expression of BMP7, phosphorylated TAK1 (p-TAK1) and total TAK1 in the cells described in panel a. **(c)** BMP7 mRNA expression levels in SW480 cells stably transfected with PPARD (PPARD^SE^) or control (Ctrl) lentivirus vector. **(d)** Expression levels of active β-catenin, BMP7, p-TAK1, TAK1, phosphorylated-p38 (p-p38), and total p38 in SW480 cells described in panel c. **(e)** Schematic map of predicted PPARD binding site (pPDBS) in the promoter of the human BMP7 gene according to Genomatix MatInspector online software. Nucleotides marked in red indicate core sequences. TSS: transcription start site. **(f)** PPARD’s binding to pPDBS in the promoter of the human BMP7 gene, measured by chromatin immunoprecipitation-quantitative PCR (-492 to -470 bp of the BMP7 promoter) in SW480 cells described in panel c. The experiments were independently replicated three times. (g) Active β-catenin expression in SW480 cells described in panel c, treated with the specific TAK1 inhibitor 5z-7 (2.5 μM) or an equal amount of control solvent (DMSO) for 24 h. **(h)** Active β-catenin expression levels in SW620 cells treated with 5z-7 (2.5 μM), PPARD antagonist GSK3787 (GSK, 1 μM), or an equal amount of DMSO for 24 h. **(i)** BMP7 mRNA expression levels in IECs of PD mice and their WT littermates. **(j)** Protein expression levels of active β-catenin, BMP7, p-TAK1, TAK1, and p38 in IECs of the mice described in panel i. **(k, l)** Representative IHC images of p-TAK1 expression **(k)** and the count of positively-stained p-TAK1 cells per field **(l)** in IECs of the mice described in panel i (n = 3 mice/group). **(m)** BMP7 mRNA expression levels in IECs of Apc^Δ580^ and Apc^Δ580^-PD littermates. **(n)** Protein expression of active β-catenin, BMP7, p-TAK1, TAK1, p-p38, and p38 in IECs of mice described in panel m. Error bars: mean ± s.e.m. \**P <* 0.05, \*\*\**P <* 0.001, **** *P <* 0.0001.

To further clarify PPARD’s mechanistic significance to the BMP7-TAK1-β-catenin signaling pathway, we employed a TAK1-specific inhibitor, 5z-7, to treat SW480 cells in which this signaling pathway was activated by PPARD overexpression. 5z-7 inhibited the increase in active β-catenin protein levels induced by PPARD overexpression in SW480 cells (Fig. 3g), thus confirming TAK1’s mechanistic contribution to PPARD activation of β-catenin. In SW620 cells with intrinsic activation of BMP7-TAK1-β-catenin signaling,(20) the specific PPARD inhibitor GSK3787 reduced active β-catenin protein levels to those achieved with 5z-7 (Fig. 3h). In vivo, PPARD increased BMP7 mRNA levels (Fig. 3i) and BMP7, p-TAK1, and p38 protein levels (Fig. 3j-l and Supplementary Fig. 2b) in IECs of PD mice. Similarly, BMP7 mRNA levels (Fig. 3m) and BMP7, p-TAK1, p38, and p-p38 protein levels were significantly higher in IECs of Apc^Δ580^-PD mice than in IECs of Apc^Δ580^ mice (Fig. 3n).

### PPARD accelerates APC mutation-driven intestinal tumorigenesis

Germline KO of PPARD in Apc^min^ mouse models produced contradictory results (11, 12). PPARD is, however, overexpressed in IECs of human CRC (6-9). We therefore sought to test the effects of PPARD overexpression in IECs of APC-mutant mice. Breeding Apc^min^ and PD mice generated Apc^min^-PD mice (Supplementary Fig. 3a), which rapidly developed a large intestinal tumor burden and required euthanasia. Apc^min^-PD mice had significantly more and larger intestinal tumors than did their littermate Apc^min^ mice (Fig. 4a,b). We also tested the effects of PPARD overexpression in IECs on tumorigenesis in Apc^Δ580^ mice, which better simulate human APC mutation-driven CRC tumorigenesis than do Apc^min^ mice(18). Apc^Δ580^-PD mice had significantly more and larger intestinal tumors than did their littermate Apc^Δ580^ mice (Fig. 4c,d and Supplementary Fig. 3b,c). In addition, Apc^Δ580^-PD mice had shorter colonic lengths (Supplementary Fig. 3d) and lower body weights (Supplementary Fig. 3e) than did Apc^Δ580^ mice. In longitudinal survival experiments, Apc^Δ580^ mice survived for longer durations (mean = 191 days) than did Apc^Δ580^-PD mice (mean = 99 days) (Fig. 4e).

**Fig 4.**
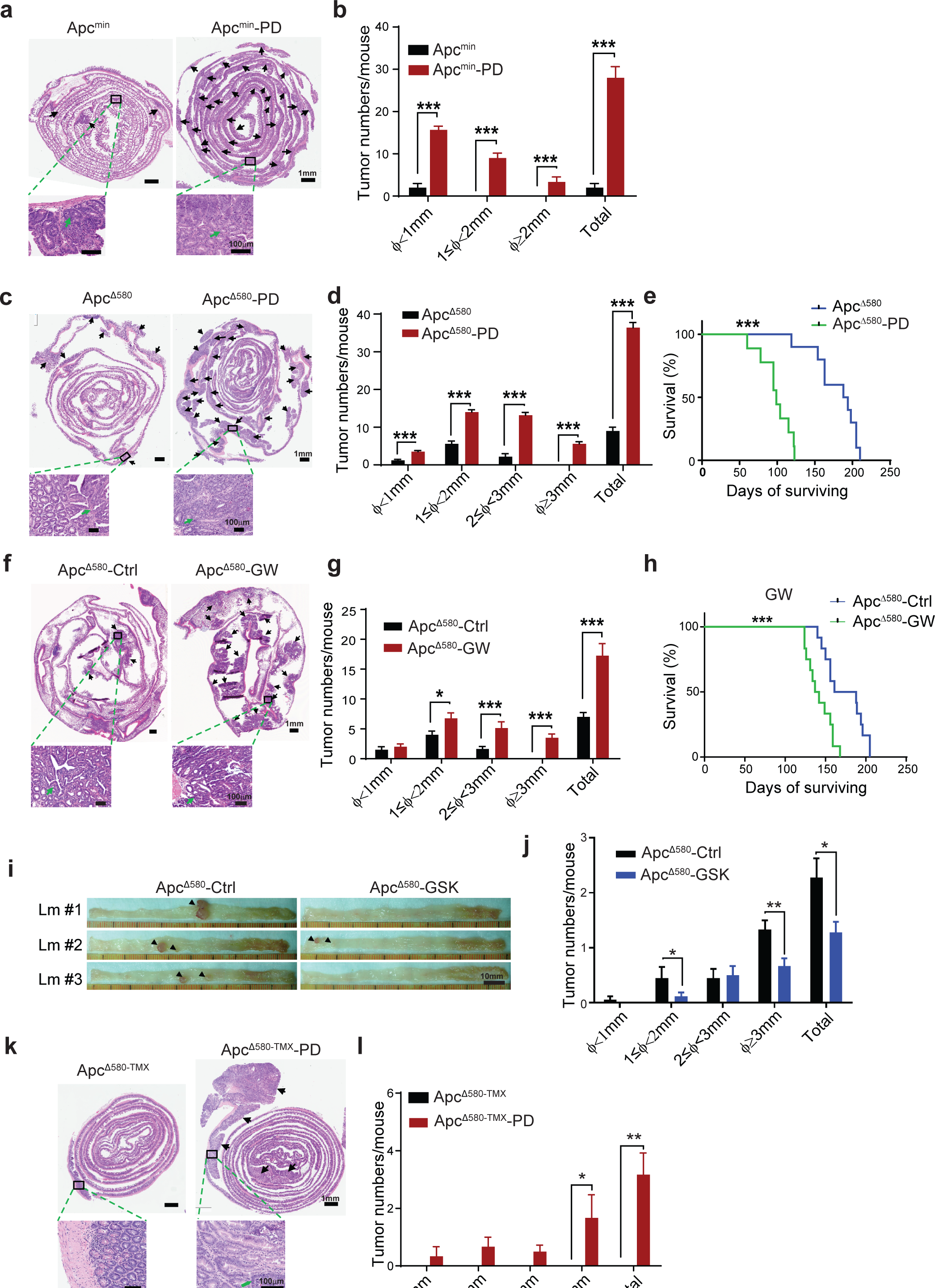
PPARD accelerates APC mutation-driven intestinal tumorigenesis in mice. **(a, b)** Effects of PPARD on intestinal tumorigenesis in Apc^min^ mice. **(a)** Representative images of H&E-stained intestines of Apc^min^ mice and Apc^min^-PD littermates at 8 weeks of age. Upper panel: Pictures of whole-length intestinal sections (Swiss roll). Arrows indicate tumors. Lower panel: higher-magnification pictures of tumor lesions (20×). **(b)** Intestinal tumor numbers and sizes for the indicated groups (n = 6 mice per group). Φ indicates tumor maximum diameter. **(c, d)** Effects of PPARD on intestinal tumorigenesis in APC^Δ580^ mice. **(c)** Representative images of H&E-stained whole-length intestinal sections of Apc^Δ580^ and Apc^Δ580^-PD mice at 14 weeks of age. Upper and lower panel picture descriptions are similar to panel a.**(d)** Intestinal tumor numbers and sizes for the indicated groups (n = 15 mice per group). (e) Survival curves of Apc^Δ580^ and Apc^Δ580^**-**PD mice (n = 20 mice per group). **(f-g)** Effects of PPARD agonist GW501516 on intestinal tumorigenesis in Apc^Δ580^ mice. **(f)** Apc^Δ580^ mice at age 4 weeks were fed a diet containing 50 mg/kg GW501516 (Apc^Δ580^-GW) or a control diet (Apc^Δ580^-Ctrl) for 10 weeks and then killed for intestinal tumorigenesis analyses. Upper and lower panel picture descriptions are similar to panel a. **(g)** Intestinal tumor numbers and sizes for the indicated groups (n = 10 mice per group). **(h)** Survival curves of indicated groups (n = 12 mice per group). **(i, j)** Effects of PPARD antagonist GSK3787 on intestinal tumorigenesis in Apc^Δ580^ mice. Apc^Δ580^ mice at age 4 weeks were fed a diet containing 200 mg/kg GSK3787 (Apc^Δ580^-GSK) or a control diet (Apc^Δ580^-Ctrl) for 12 weeks and then killed for intestinal tumorigenesis analyses. Representative colon pictures **(i)** and colonic tumor numbers and sizes **(j)** for the indicated groups (n = 18-20 mice per group). **(k, l)** Effects of PPARD on intestinal tumorigenesis driven by adult-onset intestinally targeted APC mutation. Apc^Δ580^ mutation was induced in the mice at age 6 weeks via tamoxifen-controlled Cre-recombinase expression driven by CDX-2 promoter (Apc^Δ580-TMX^). Apc^Δ580-TMX^ mice without or with intestinal PPARD overexpression (Apc^Δ580-TMX^-PD) were followed for 55 weeks before they were killed for intestinal tumorigenesis analyses. **(k)** Upper and lower panel picture descriptions are similar to panel a. **(l)** Intestinal tumor numbers and sizes for the indicated groups (n = 8 mice per group). Error bars: mean ± s.e.m. \**P <* 0.05, \*\**P <* 0.01, \*\*\**P <* 0.001.

We next tested the effects of the PPARD agonist GW501516 in Apc^Δ580^ mice using a diet containing 50 mg/kg GW501516. GW501516 diet produced significantly more and larger intestinal tumors than did the control diet (Fig. 4f,g and Supplementary Fig. 3f,g). The median survival of GW50106-treated Apc^Δ580^ mice was shorter (mean = 140 days) than that of Apc^Δ580^ mice fed a control diet (mean = 191 days) (Fig. 4h). In complementary testing, Apc^Δ580^ mice fed a diet containing 200 mg/kg GSK3787 (a highly specific PPARD antagonist (21)) had significantly fewer and smaller colonic tumors than did mice fed a control diet (Fig. 4i, j). In particular, the number of tumors larger than 3 mm decreased from 1.33 tumors per control-treated mouse (95% confidence interval [CI]: 0.99-1.67) to 0.67 tumors per GSK3787-treated mouse (95% CI: 0.37-0.96) (Fig. 4j).

We also examined the effects of PPARD overexpression on adult-onset APC mutation using tamoxifen-inducible Apc^Δ580^ mice (Apc^Δ580-TMX^) to simulate the most common form of human CRC (i.e., sporadic) with an adult-onset APC mutation (Supplementary Fig. 3a, Row #3). Apc^Δ580-TMX^ mice with heterozygote APC mutations induced at 5 weeks of age had no visible intestinal tumors (only 1 small adenoma was detected microscopically), even after 55 weeks of follow-up. In contrast, their littermates with PPARD overexpression in IECs (Apc^Δ580-TMX^-PD) developed visible tumors, including large CRCs (*P* = 0.025) (Fig. 4k,l and Supplementary Fig. 3h).

### PPARD promotes CRC invasiveness

The transformation to invasive CRC occurs in large adenomas (3). PPARD overexpression in IECs significantly increased the number of large tumors in Apc^min^-PD, Apc^Δ580^-PD, and Apc^Δ580-TMX^-PD mice and in GW501516-treated Apc^Δ580^ mice (Fig. 4b,d,g,j,l). We found invasive intestinal tumors in 86% of Apc^Δ580^-PD mice but only 16% of their littermate Apc^Δ580^ mice (*P* = 0.029) (Fig. 5a,b). The mean number of invasive tumors per mouse increased from 0.14 (95% CI: 0.2-0.49) in Apc^Δ580^ mice to 2.29 (95% CI: 1-3.56) in Apc^Δ580^-PD mice (*P* = 0.0056) (Fig. 5b). When APC mutation was delayed until the mice reached adulthood, invasive tumors developed in Apc^Δ580-TMX^-PD mice but not in their Apc^Δ580-TMX^ littermates (Fig. 5c). To assess the clinical relevance of these findings, we compared PPARD expression in paired samples of human colorectal adenomas and the centers and fronts of invasive tumors (N = 41 patients). Immunohistochemistry composite expression scores of PPARD expression were significantly higher in the CRC invasive fronts than in their paired tumor centers or adenomas (Fig. 5d,e).

**Fig 5.**
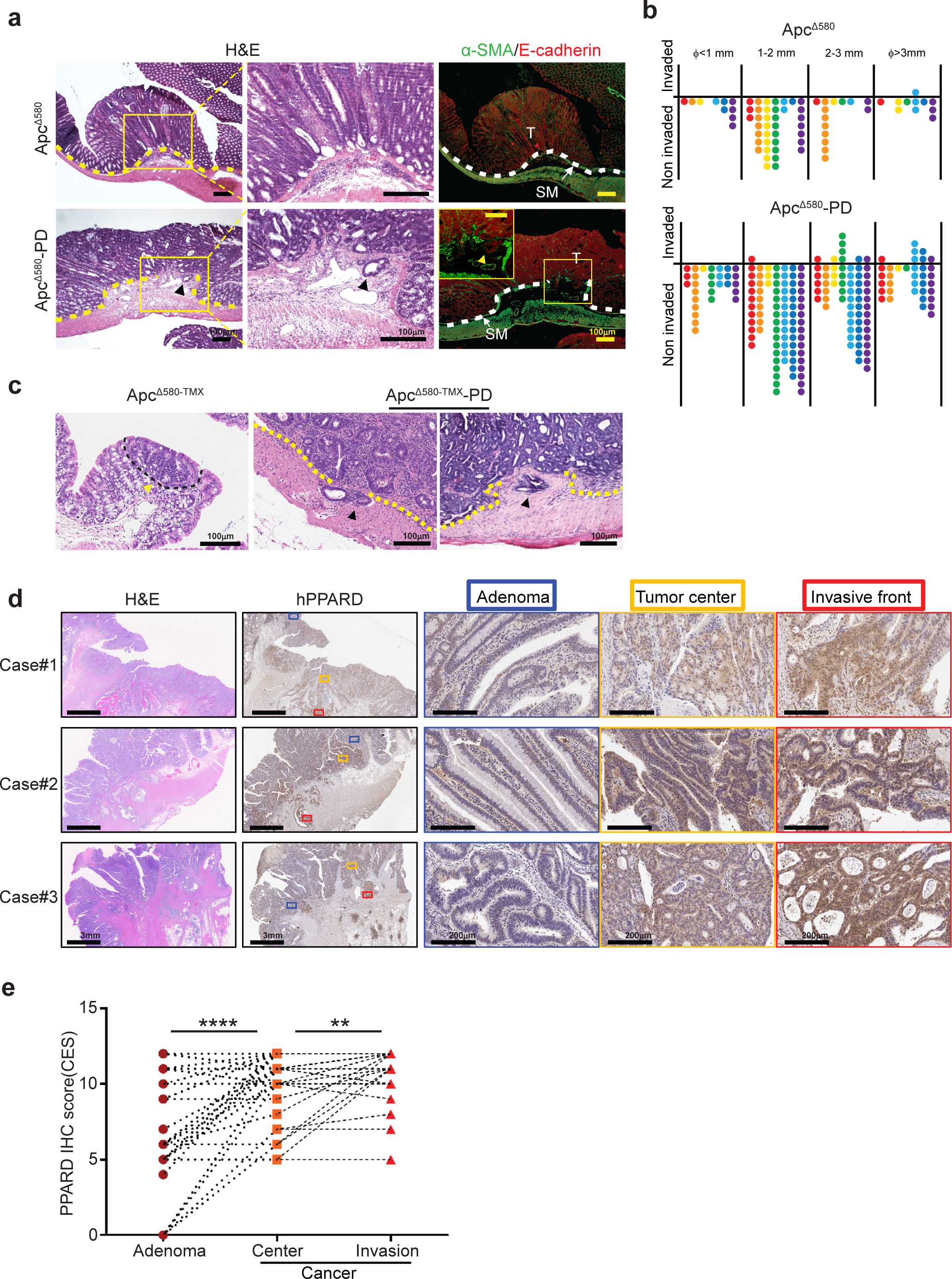
PPARD promotes intestinal tumor invasion. **(a, b)** Intestinal PPARD overexpression promotes invasion of intestinal tumors in Apc^Δ580^ mice. Intestines of Apc^Δ580^ and Apc^Δ580^-PD littermate mice (20 weeks old) were evaluated for tumor invasiveness. **(a)** Representative intestinal section photographs of noninvasive adenoma (upper) and invasive adenocarcinoma (lower) in the indicated mouse groups. Pictures of low (left) and high (middle) magnifications of H&E staining and immunofluorescence staining (right) for E-cadherin (red) and α-SMA (green). T, tumor; SM, submucosa. Arrowheads indicate submucosal invasion of cancer cells. **(b)** Numbers of invasive and noninvasive tumors per mouse for the indicated groups, scored in whole-mount intestinal (Swiss roll) sections (n = 7 mice/group). **(c, d)** Apc^Δ580^ mutation was induced by tamoxifen treatment of littermate mice without (Apc^Δ580-TMX^) or with intestinal PPARD overexpression (Apc^Δ580-TMX^-PD) at 6 weeks of age. Mice were followed for 55 weeks prior to being killed to assess intestinal tumor invasiveness. **(c)** H&E staining photographs showing the only microscopically identified adenoma in the Apc^Δ580-TMX^ group and 2 representative invasive adenocarcinomas in the Apc^Δ580-TMX^-PD group. **(d)** Numbers of invasive and noninvasive tumors per mouse for the indicated groups were scored in whole-mount intestinal sections (n = 5-6 mice/group). **(e,f)** PPARD expression in human colorectal cancer invasive front. **(e)** Representative PPARD IHC staining images of human paired colon adenomas, colorectal cancer centers (tumor center), and tumor invasive fronts of 3 patients. **(f)** Total composite expression score (CES) of nucleus and cytoplasm IHC staining of PPARD for the paired adenomas, colorectal cancer centers, and invasive fronts as described in panel e (n = 41 patients). Error bars: mean ± s.e.m. \*\**P <* 0.01, \*\*\*\**P <* 0.0001.

### PPARD regulates PDGFRβ expression in CRC tumorigenesis

We screened for downstream target genes of PPARD that promoted intestinal tumorigenesis using comparative functional proteomics reverse phase protein array analyses of IECs from Apc^Δ580^ and Apc^Δ580^-PD mice. These analyses showed different proteomic patterns for the Apc^Δ580^-PD and Apc^Δ580^ mice (Fig. 6a). Volcano plot analyses identified 7 differentially expressed proteins with at least a 2-fold expression change and 2-sided *t*-test *P* values of less than .05. Six of these proteins were upregulated (connexin 43, AKT serine/threonine kinase 1 [AKT1], platelet-derived growth factor receptor β [PDGFRβ], cyclin-dependent kinase 1 [CDK1], eukaryotic translation initiation factor 4 γ [EIF4G], and phosphorylated ribosomal protein S6 [rpS6] at residues 235/236 and 240/244), while only 1 protein (caspase 7-cleaved) was downregulated (Fig. 6b). Independent experiments confirmed PPARD upregulation of PDGFRβ protein and mRNA expression in Apc^Δ580^-PD mice (Fig. 6c,d) and PD mice (Fig. 6e,f). In subsequent clinical relevance assessments using paired normal and CRC tissue samples from MD Anderson colorectal cancer patients (n = 22), PPARD and PDGFRβ mRNA levels were concomitantly higher in CRC tissues than in paired normal tissues in 20 of 22 patients. Relative mRNA level (cancer vs. normal) ratios were above 2 for PDGFRβ (mean ± SEM: 33.76 ± 15.5; 95% CI: 2.46-65.05; *P* < 0.0001) and PPARD (8.88 ± 2.4; 95% CI: 3.9-13.87; *P* < 0.0001) (Fig. 6g,h). In data from The Cancer Genome Atlas (TCGA) provisional colorectal cancer database, PDGFRβ and PPARD mRNA levels were significantly correlated, with a tendency towards co-occurrence (log odds ratio = 2.348; *P* < 0.001) (Fig. 6i). On examining PDGFRβ’s mechanistic significance to PPARD’s promotion of CRC invasion, we found that PPARD overexpression in CT26 mouse CRC cells significantly increased cell migration and that DMPQ (a specific PDGFRβ inhibitor(22)) reduced cell migration promoted by PPARD (Fig. 6j).

**Fig 6.**
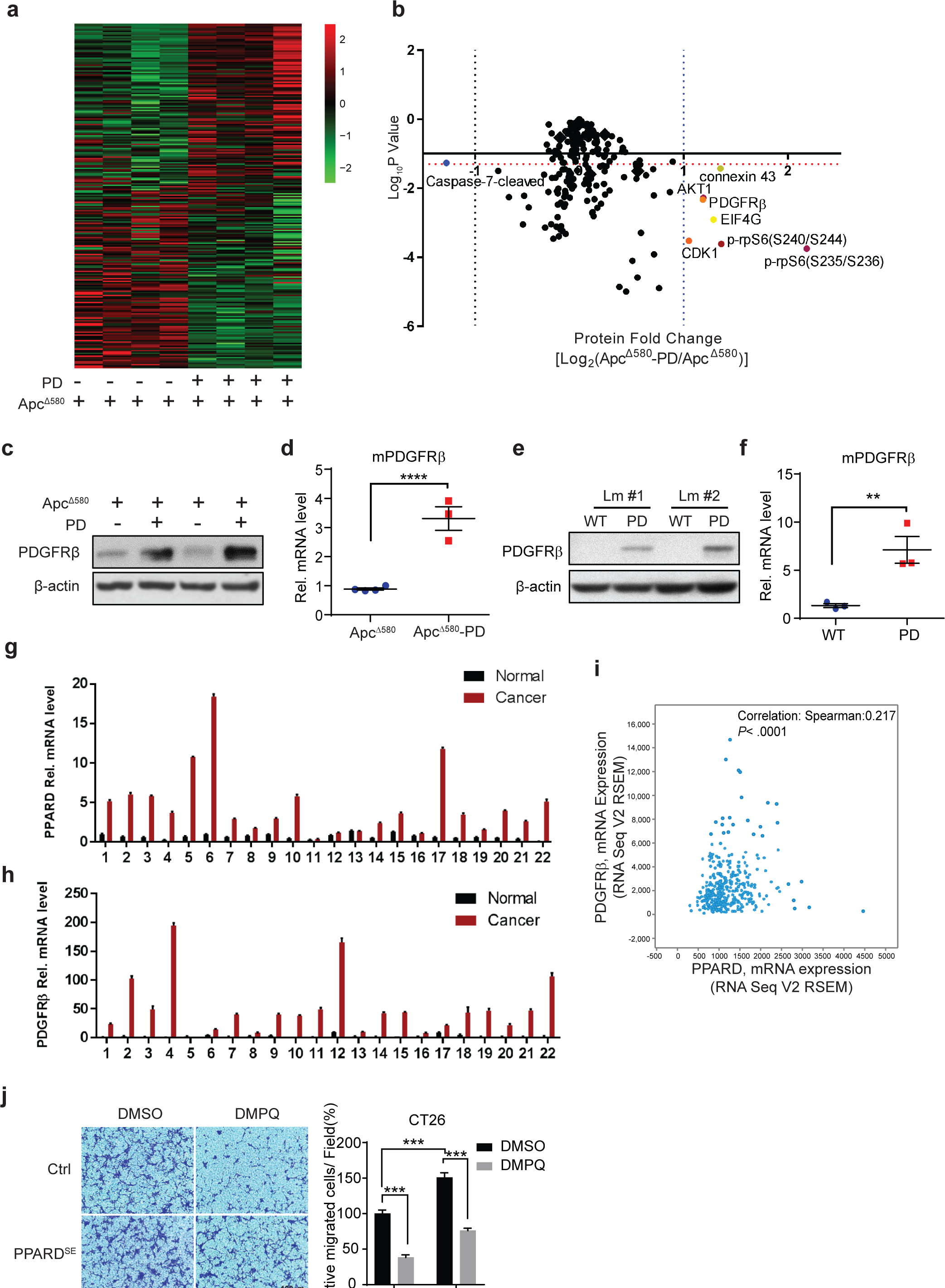
PPARD transcriptionally upregulates PDGFRβ expression in mice with or without APC mutation. **(a, b)** RPPA analyses were performed on normal intestinal epithelial cells (IECs) of Apc^Δ580^ and Apc^Δ580^-PD littermate mice at 10 weeks of age (n = 4 mice per group). Results are presented as a heat map **(a)** and a volcano plot of the differentially expressed genes **(b)**. The horizontal orange dotted line indicates the threshold, *P* = 0.05. The vertical blue dotted line on the right indicates where the log_2_ ratio was > 1, and the black vertical dotted line on the left indicates where the log_2_ ratio was < -1. **(c-f)** Effects of PPARD overexpression on PDGFRβ protein and mRNA expression levels in IECs of APC ^Δ580^ and APC ^Δ580^-PD **(c,d)** and PD and WT littermate (Lm) mice **(e,f)**. **(g, h)** PPARD **(g)** and PDGFRβ **(h)** mRNA levels in paired patient colorectal cancer and normal mucosa samples (n = 22). **(i)** Spearman correlation analysis of PPARD and PDGFRβ mRNA expression in the TCGA Provisional Colorectal Adenocarcinoma database (n = 633). **(j)** Effects of PDGFRβ inhibitor DMPQ on PPARD promotion of colon cancer cell migration. CT26 mouse colon cancer cells were stably transfected with a mouse PPARD (PPARD^SE^) or control (Ctrl) lentiviral vector were treated with DMPQ (1 μM) or vehicle solvent (DMSO) for 36 h. Left: representative photomicrographs of migrated cells. Right: migrated cells in at least 4 random individual fields per insert membrane were counted. The experiments were independently replicated three times. Error bars: mean ± s.e.m. ** *P <* 0.01, *** *P <* 0.01, **** *P <* 0.0001.

### PPARD activates AKT-rpS6 signaling to promote intestinal tumorigenesis

Independent experiments confirmed that PPARD upregulated AKT1 protein expression in PD (Fig. 7a) and Apc^Δ580^-PD mice (Fig. 7b). In contrast, AKT2 expression remained unchanged (Fig. 7a,b and Supplementary Fig. 4a,b). PPARD also increased AKT1 phosphorylation and activation as measured by phosphorylation of downstream target proteins rpS6 and glycogen synthase kinase 3β (GSK3β) (Fig. 7a,b and Supplementary Fig. 4c,d). Furthermore, we found that PPARD overexpression upregulated AKT1 mRNA levels in IECs of PD and Apc^Δ580^-PD mice (Fig. 7c,d). When we integrated data from the TCGA colorectal cancer databases to determine the relevance of these results to human CRC, we found significant correlation between PPARD and AKT1 mRNA expression levels (Fig. 7e). Furthermore, PPARD overexpression via lentivirus transduction increased AKT1 mRNA and protein expression and phosphorylation (Fig. 7f,g). PPARD KO HCT-116 cells (KO1) had lower AKT1 and p-rpS6 expression levels than did parental HCT-116 WT cells (Supplementary Fig. 4e). PPARD downregulation using 2 independent PPARD small interfering RNAs (siRNAs) in SW480 cells reduced AKT1 mRNA and protein levels and phosphorylation (Fig. 7h,i). Finally, we evaluated whether PPARD as a transcriptional factor acted directly by binding to the AKT1 promoter by identifying a pPDBS in an in silico search (Fig. 7j). PPARD overexpression in SW480 cells via PPARD lentivirus transduction significantly increased PPARD binding activity at the identified pPDBS site of the AKT1 promoter (Fig. 7k).

**Fig 7.**
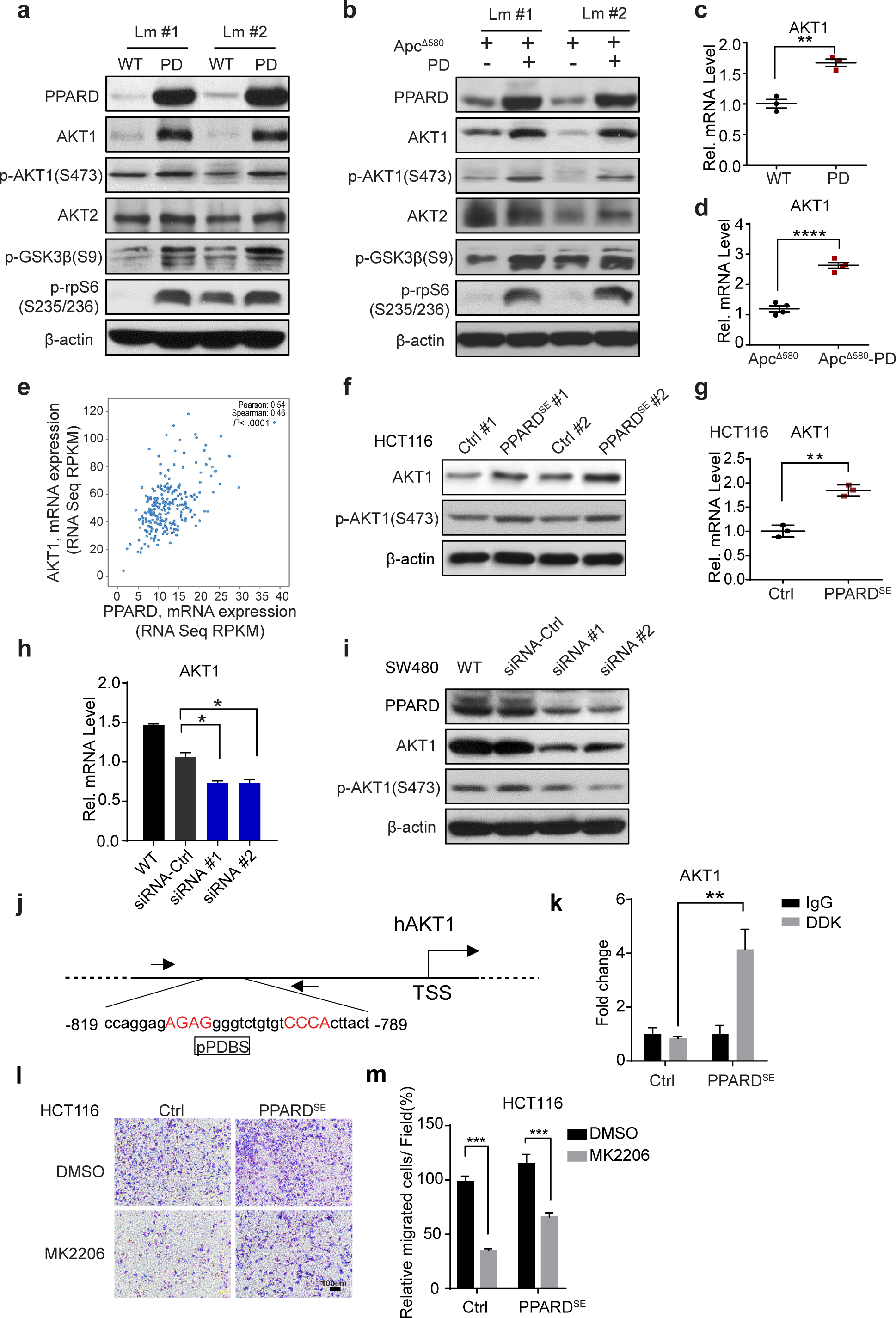
PPARD transcriptionally activates AKT1 pathway in intestinal epithelial cells. (**a, b**) Effects of PPARD on AKT signaling pathway in intestinal epithelial cells (IECs). Protein expression levels of AKT1, p-AKT1(S473), AKT2, p-GSKβ, and p-rpS6(S235/236) in IECs of PD mice **(a)**, Apc^Δ580^-PD mice **(b)**, and their corresponding control littermates (Lm). **(c, d)** AKT1 mRNA expression levels in PD **(c)** and Apc^Δ580^-PD mice **(d)** and their control littermate mice. **(e)** Correlation analysis of PPARD and AKT1 mRNA expression in the TCGA Colorectal Adenocarcinoma database (*Nature*, 2012) database (n = 244). **(f)** Protein expression levels of AKT1 and p-AKT1(S473) in HCT-116 cells stably transfected with a PPARD plasmid (PPARD^SE^) or a control (Ctrl) plasmid. **(g)** AKT1 mRNA levels in the HCT-116 cells described in panel f. **(h)** AKT1 mRNA levels in SW480 cells transfected with 2 independent PPARD siRNAs or a control siRNA. **(I)** AKT1 and p-AKT1(S473) protein expression levels in the SW480 cells transfected with PPARD siRNA described in panel h. **(j)** Schematic map of the predicted PPARD binding site (pPDBS) in the promoter region of human AKT1 (hAKT1) according to Genomatix MatInspector online software. Nucleotides in red indicate core sequences. TSS: transcription start site. **(k)** PPARD binding to the pPDBS of the AKT1 promoter shown in panel j, measured by a chromatin immunoprecipitation-quantitative PCR assay in HCT-116 cells with or without PPARD overexpression, as described in panel f. **(l,m)** Effects of AKT1 inhibition on PPARD promotion of colon cancer cell migration. Migration was assessed in HCT-116-PPARD^SE^ and HCT-116-Ctrl cells treated with an AKT inhibitor (MK2206) (1 μM) for 36 h. **(l)** Representative photomicrographs of migrated cells for the indicated treatments and PPARD overexpression conditions. **(m)** Quantitative data for the experiments described in panel l (at least 4 random individual fields per insert membrane were counted). The experiments were independently replicated three times. Error bars: mean ± s.e.m. \**P* < 0.05, \*\**P* < 0.01, \*\*\**P* < 0.001, \*\*\*\**P* < 0.0001.

We examined the biological significance of AKT upregulation by PPARD for CRC invasiveness by conducting a cell migration assay using an AKT-selective inhibitor, MK2206. PPARD overexpression in HCT-116 cells significantly increased the number of migrated cells, and MK2206 attenuated these effects (Fig. 7l,m).

### PPARD increases protein synthesis to promote intestinal tumorigenesis

In further independent reverse phase protein array experiments, we confirmed that CDK1 expression was significantly higher in PD and Apc^Δ580^-PD mice than in their control littermates at both the mRNA (Fig. 8a,b) and protein (Fig. 8c,d) levels. Similarly, EIF4G protein expression was higher in PD and Apc^Δ580^-PD mice than in their Apc^Δ580^ littermates (Fig. 8c,d). Because EIF4G is critical to the initiation of protein translation,(23) we investigated whether intestinal PPARD overexpression increased protein translation by measuring ribosomal RNA. Intestinal PPARD overexpression markedly increased ribosomal RNA levels in both PD and Apc^Δ580^-PD mice compared to their control littermates (Fig. 8e,f).

**Fig 8.**
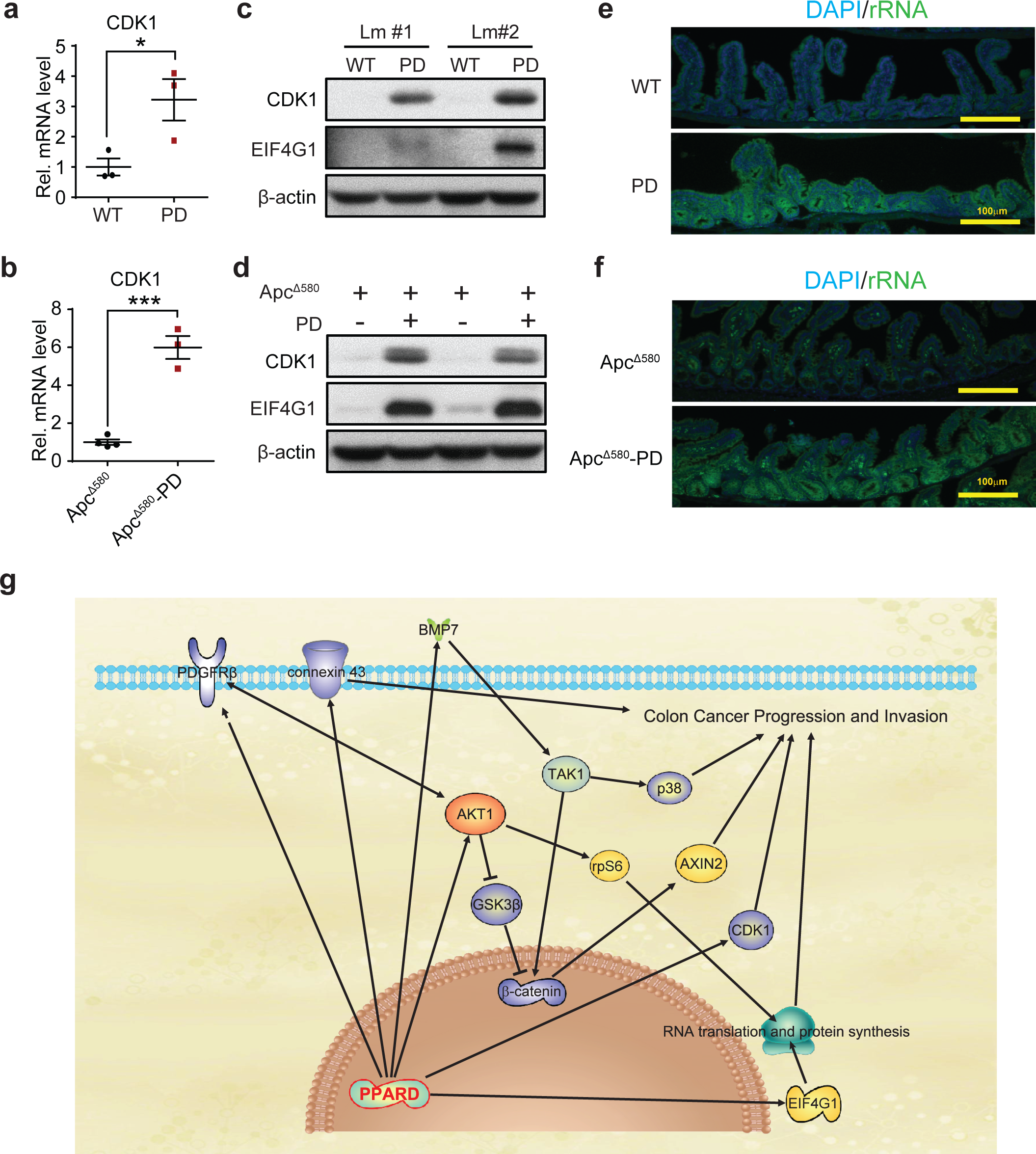
PPARD promotes cell cycle and protein synthesis. **(a, b)** CDK1 mRNA expression levels in intestinal epithelial cells (IECs) of PD mice **(a)**, Apc^Δ580^-PD mice **(b)**, and their corresponding control littermates (Lm). **(c, d)** CDK1 and EIF4G1 protein expression levels in IECs from PD **(c)** and Apc^Δ580^-PD **(d)** mice and their corresponding control littermates. **(e,f)** Immunofluorescence staining of rRNA using a Y10b antibody (Green) for the intestinal tissues from PD **(e)** and Apc^Δ580^-PD mice **(f)** and their corresponding control littermates. **(g)** Conceptual scheme of proinvasive pathways that are regulated by PPARD to promote colorectal cancer progression and invasion. The solid lines: direct regulation; the dotted lines: indirect regulation. Error bars: mean ± s.e.m. \**P* < 0.05, \*\*\**P* < 0.001.

## DISCUSSION

We found that PPARD potentiated β-catenin activation in IECs via upregulation of BMP7/TAK1 signaling and promoted colorectal tumorigenesis progression and invasion by upregulating multiple pro-invasive pathways, including PDGFRβ, AKT1, CDK1, EIF4G, and connexin 43(Fig. 8g).

Our new data provide important new insights that address the controversy regarding PPARD’s effects on β-catenin promotion of CRC. Using in vivo and in vitro models with genetic and pharmacological modulation of PPARD expression and activation, we clearly demonstrated that PPARD increased β-catenin activation downstream of APC mutations. In these models, PPARD activated β-catenin and upregulated its target genes, including c-Myc, which enhances progenitor colonic crypt cell proliferation to promote CRC (24). Indeed, PPARD upregulation in mouse IECs dramatically increased intestinal progenitor cell self-renewal, especially in the form of immature spheroid organoids. Activation of PPARD via high-fat diet or pharmaceutical ligands has been reported to enhance intestinal organoid formation in cells of Apc^min^ mice(15). Our findings, however, demonstrate for the first time that PPARD overexpression in APC-WT IECs is sufficient to enhance intestinal progenitor cell self-renewal in the absence of APC mutations. These data address the issues of ligand off-target effects and whether PPARD’s effects depend on APC mutations. Our finding that PPARD overexpression in IECs increased β-catenin activation both in APC-WT mice and in APC-WT HCT-116 cells also indicates that these effects of PPARD are downstream of APC.

We found that PPARD activates β-catenin via upregulating BMP7 to phosphorylate TAK1. Our data demonstrate for the first time that PPARD transcriptionally upregulates BMP7 expression, which subsequently activates TAK1/β-catenin signaling. Previously, the BMP7/TAK1/β-catenin signaling pathway was reported to be selectively active in APC- and KRAS-mutant colon cancer cells and to contribute to therapeutic resistance(20). We showed that a PPARD inhibitor, GSK3787, suppressed β-catenin activation in APC- and KRAS-mutant SW620 colon cancer cells. More importantly, we demonstrated for the first time that PPARD upregulation activated BMP7/TAK1/β-catenin signaling not only in colon cancer cell lines with both APC and KRAS mutations (i.e., SW480 cells) or with WT APC and mutant KRAS (i.e., HCT-116 cells), but also in vivo in mice with and without APC mutations (PD and Apc^Δ580^-PD). These data clearly demonstrate that PPARD regulates this therapeutic resistance pathway.

In addition, we determined that PPARD transcriptionally upregulates BMP7 to activate TAK1. Prior reports have been inconsistent regarding PPARD’s relation to TAK1. PPARD agonists have been reported to inhibit TAK1 phosphorylation in rodent renal tubular (25) and peritoneal mesothelial cells (26). However, the concentrations of PPARD agonists in these experiments were above the specific PPARD activation range and PPARD downregulation or overexpression failed to alter the agonist’s effects (25), thus suggesting that non-PPARD-mediated mechanisms are involved. Other researchers have questioned whether PPARD affects TAK1 activity because PPARD downregulation via siRNA in HeLa cells reduced TAK1 phosphorylation, whereas PPARD overexpression in HEK293T cells failed to increase TAK1 phosphorylation (27). In contrast with these ambiguous results, our data clearly show that PPARD increases TAK1 phosphorylation in both in vitro and in vivo PPARD gain- and loss-of-function models. Moreover, our results elucidate the mechanism by which PPARD regulates TAK1: transcriptional upregulation of BMP7. BMP7, which is upregulated or amplified in 16% to 24% of human colorectal cancers in the TCGA databases (Supplementary Fig. 5a), negatively affects CRC patients’ survival (Fig. 5d,e and Supplementary Fig. 5c) and promotes colon cancer invasiveness (28). TAK1 phosphorylation is being investigated clinically as a molecular target in the treatment of human cancers (29). Our identification of PPARD, a druggable protein, as a regulator of the BMP7/TAK1 signaling pathway suggests a novel approach to therapeutically target BMP7/TAK1.

Our findings also provide important new insights regarding PPARD’s effects on APC mutation-driven CRC tumorigenesis. Our novel mouse models of PPARD overexpression in IECs with APC mutations clearly show that PPARD strongly promotes CRC tumorigenesis. Prior studies were limited to PPARD KO modeling in APC-mutant mice. PPARD is, however, upregulated (6-9), not deleted, in human CRC (Supplementary Fig. 5d). Thus, we tested PPARD’s effects not only in Apc^min^ mice, which develop small intestinal adenomas, but also in Apc^Δ580^ mice, in which intestinally targeted APC mutations better simulate human CRC. In contrast with the conflicting data from experiments with Apc^min^ mice with germline PPARD KO, in all of our models, PPARD strongly potentiated APC mutation-driven CRC tumorigenesis. Other researchers have reported that PPARD activation with GW501516 significantly increased tumor numbers and sizes in the small intestines in Apc^min^ mice(30). Our data showed not only that GW501516 significantly increased intestinal tumorigenesis in large intestines of Apc^Δ580^ mice but also, for the first time, that a PPARD inhibitor, GSK3787, had antitumorigenic activity in in vivo CRC models. APC mutations occur later in life in the majority of patients with CRC; fewer than 1% have germline APC mutations(31). Our novel Apc^Δ580-TMX^ mouse model, in which the onset of APC mutation is delayed to early adulthood to better simulate the most common human CRC pattern, showed even more striking protumorigenic effects of PPARD overexpression in IECs. Thus, our extensive and in-depth modeling of intestinal PPARD in relation to APC mutations clearly establishes PPARD’s role in promoting CRC tumorigenesis.

APC mutations alone are insufficient to induce CRC invasiveness (3, 32). Our findings demonstrate the mechanistic significance of PPARD overexpression in IECs for CRC invasiveness. PPARD overexpression in intestinal cells significantly increased CRC invasiveness in both our early- and late-onset APC mutation mouse models. These novel findings are relevant to human CRC; using CRC patient samples, we found that PPARD was more strongly upregulated in CRC invasive fronts (an indicator of CRC invasiveness in humans (33, 34)) than in paired adenomas or even invasive cancer centers.

Furthermore, functional proteomic analyses identified multiple pathways, including PDGFRβ, AKT1, CDK1, EIF4G, and connexin 43, that strongly promote CRC tumorigenesis and invasiveness. PPARD upregulated connexin 43 in IECs in vivo, which is in agreement with our previously published data from differential transcriptome analyses of HCT-116 and KO1 cells (9). Our results however demonstrate for the first time that PDGFRβ, a key gene in colorectal adenoma-to-carcinoma progression(35), is upregulated by PPARD to promote CRC tumorigenesis. The AKT/mTOR signaling pathway integrates various upstream oncogenic effector signals (e.g., PDGFRs) to modulate downstream targets (e.g., ribosomal protein S6 kinase 1 [S6K1], EIF4E binding proteins) and promote tumor progression and invasiveness (36). PPARD has been reported to activate AKT in mouse keratinocytes (37), A549 non-small cell lung cancer cells (38), and mammary epithelial cells (39). Our data demonstrate for the first time that: (1) PPARD specifically upregulates AKT1, but not AKT2, expression and phosphorylation in in vitro and in vivo CRC tumorigenesis models; (2) PPARD transcriptionally regulates AKT1; and (3) PPARD is strongly correlated with AKT1 expression in human CRC. In addition, PPARD increased GSK3β phosphorylation, which enhances various protumorigenic mechanisms, especially aberrant β-catenin activation(40). PPARD agonists have been reported to increase AKT and GSK3β phosphorylation in rodent hearts challenged by ischemia or sepsis (41, 42). Our data, however, show for the first time that PPARD increases GSK3β phosphorylation to drive colorectal tumorigenesis.

Dysregulation of mRNA translation plays a very important role in tumorigenesis and cancer progression(43). Tumorigenesis enhances aberrant mRNA translation via multiple mechanisms, including AKT/mTOR activating phosphorylation of S6K1/2 and inactivating phosphorylation of EIF4E-binding proteins to release EIF4E, which subsequently forms a protein complex with other eukaryotic translation initiation factors (e.g., EIF4G) and initiates mRNA translation(43). Enhancement of the EIF4E-EIF4G interaction promotes the development of cancer-cell resistance to BRAF and MEK inhibitors (44), and EIF4G overexpression in solid tumors promotes tumor progression(43). Our novel finding that PPARD upregulated EIF4G expression in IECs is an additional mechanism by which PPARD promotes tumorigenesis and progression. We also demonstrated for the first time that PPARD upregulated CDK1 in IECs. CDK1 regulates cell cycle-dependent (e.g., chromosome segregation, DNA repair(45)) and -independent functions (e.g., promotion of protein translation independent of AKT (46)) to promote tumorigenesis and progression and therefore is considered a potential cancer therapeutic target(47). In sum, these novel findings demonstrate PPARD’s pleiotropic effects in IECs to enhance RNA translation and cancer invasiveness.

In conclusion, our findings demonstrate that PPARD strongly accelerates APC mutation-driven colorectal tumorigenesis and tumor invasion via multiple important protumorigenic pathways, including BMP7/TAK1/β-catenin, PDGFRβ, AKT1, EIF4G, CDK1, and connexin 43 (Fig. 8g). These findings establish PPARD’s pivotal role in promoting CRC progression and its potential as a preventive and therapeutic target.

## METHODS

### Mouse models

Mouse care and experimental protocols were approved by and conducted in accordance with the guidelines of the Animal Care and Use Committee of The University of Texas MD Anderson Cancer Center. We generated mice with targeted PPARD overexpression in IECs (PD mice) via a villin promoter as described previously (48). C57BL/6J-*Apc*^*Min*^/J (Apc^min**/+**^, stock #002020), B6.Cg-Tg (CDX2-Cre)101Erf/J (CDX2-Cre, stock # 009350), and B6.Cg-Tg(CDX2-Cre/ERT2)752Erf/J (CDX2-CreERT2, stock #022390) mice were purchased from Jackson Laboratory. Apc^Δ580^ floxed (Apc^Δ580^-flox) mice, in which APC exon 14 is flanked with loxp sites, were a gift from Dr. Kenneth E. Hung(49). Breeding Apc^Δ580^-flox mice with Cre recombinase-expressing (CDX2-cre or CDX2-CreERT2) mice deleted APC exon 14 and consequently generated a codon 580 frame-shift mutation without (Apc^Δ580^ mice) or with tamoxifen treatment (Apc^Δ580-TMX^ mice) (49).

### Cell lines

The human colon cancer cell lines SW480 and SW620 and the mouse colon cancer cell line CT26 were purchased from ATCC; HCT-116 parental and peroxisome proliferator-activated receptor-δ (PPARD)-knockout (KO1) cells were kindly provided by Dr. Bert Vogelstein (Johns Hopkins University, Baltimore, MD). HCT-116 and KO1 cells were authenticated by short tandem repeat analyses by the Characterized Cell Line Core Facility at The University of Texas MD Anderson Cancer Center. HCT-116 and KO1 cells were cultured in McCoy’s 5A medium. SW480, SW620, and CT26 cells were cultured in RPMI-1640 medium supplemented with 10% fetal bovine serum (FBS) (VWR International). All culture media were supplemented with 10% FBS, 2 mM L-glutamine, and 1% penicillin/streptomycin (Life Technologies).

### Clinical samples

De-identified sections from paraffin-embedded tissue blocks of archived surgical pathology materials were obtained from the colorectal tumor tissue repository at MD Anderson Cancer Center with Institutional Review Board (IRB) approval. These sections contained areas of adenomatous polyps and cancer arising within the polyps and paired normal-appearing colonic mucosa from at least 5 cm from the tumor edge. Residual surgical pathology samples were obtained from 41 colon cancer patients who underwent surgical resection of primary colon cancer without prior exposure to chemotherapy or radiation therapy and consented to the future use of their residual tissue for research. An experienced colon pathologist (R.B.) confirmed that these sections contained colonic adenoma and colon cancer with the tumor center and its invasive front in the same hematoxylin and eosin (H&E)-stained section for each case.

RNA samples from paired normal and malignant colonic mucosa from patients with stage III colon cancer were obtained with IRB approval as described previously.(9)

### Mouse intestinal tumorigenesis testing

#### Apc^min^ mice with or without PPARD overexpression in intestinal epithelial cells

Apc^min^ mice were bred with villin-PPARD mice (PD mice) to generate Apc^min^;PPARD (Apc^min^-PD) mice. Apc^min^ and Apc^min^-PD mice (n = 6 mice per group) were followed to the age of 8 weeks and then euthanized and necropsied.

#### Apc^Δ580^ mice with or without specific PPARD overexpression in the intestines

Apc^Δ580^ mice were bred with PD mice to generate double-transgene Apc^Δ580^;PPARD (Apc^Δ580^-PD) mice. Apc^Δ580^ and Apc^Δ580^-PD mice were followed until the age of either 14 weeks (n = 15 mice per group) or 22 weeks (n = 7 mice per group) and then euthanized to evaluate intestinal tumorigenesis.

#### Treatment of Apc^Δ580^ mice with PPARD agonist GW501516

Apc^Δ580^ mice at age 4 weeks were fed a diet containing 50 mg/kg GW501516 or the same diet without GW501516 (control diet) (Envigo) for 10 consecutive weeks (n = 10 mice per group). The mice were then euthanized and examined for intestinal tumorigenesis. The chemical GW501516 was synthesized by the Translational Chemistry Core Facility at MD Anderson Cancer Center, and its authenticity was confirmed by liquid chromatography-mass spectrometry standard GW501516 purchased from an independent source (Cat# SML1491, Sigma).

#### Treatment of Apc^Δ580^ mice with PPARD antagonist GSK3787

Apc^Δ580^ mice at age 4 weeks were fed a diet containing 200 mg/kg GSK3787 or the same diet without GSK3787 (control diet) (Envigo) for 12 consecutive weeks (n = 18-20 mice per group), then euthanized and examined for intestinal tumorigenesis. GSK3787 was synthesized by the Applied Cancer Science Institute at MD Anderson Cancer Center, and its authenticity was confirmed by liquid chromatography-mass spectrometry analyses using standard GSK3787 purchased from Sigma (Cat# G7423).

#### Apc^Δ580-TMX^ mice with or without specific PPARD overexpression in the intestines

Apc^Δ580^**-^TMX^** mice (generated by breeding Apc^Δ580^-flox with CDX2-CreERT2) were bred with PD mice to generate Apc^Δ580^**-^TMX^**;PPARD (Apc^Δ580^**-^TMX^**-PD) mice. Apc^Δ580^**-^TMX^** and Apc^Δ580^**-^TMX^**-PD mice (n = 8 mice per group) were treated with tamoxifen (Sigma) dissolved in corn oil (0.75 mg/10 g mice) by gavage once a day for 3 consecutive days and then followed for up to 55 weeks before being euthanized and examined for intestinal tumorigenesis.

### Mouse tumorigenesis survival experiments

Two independent mouse intestinal tumorigenesis survival experiments were performed for the following groups: (1) Apc^Δ580^ and Apc^Δ580^-PD mice (n = 20 mice per group) and (2) Apc^Δ580^ mice on a diet containing 50 mg/kg GW501516 or a control diet (n = 12 mice per group). For each experiment, mice were followed until they required euthanasia on the basis of 1 of the following preset criteria: (1) persistent rectal bleeding for 3 consecutive days or (2) weight loss of more than 20%.

### Mouse intestinal tumorigenesis evaluation

The mice were euthanized, and their abdomens were opened. The intestines from the rectum to the base of the stomach were removed and washed with phosphate-buffered saline (PBS). For whole intestinal mount (Swiss roll) analyses, one end of the intestine was ligated, and the intestine was inflated with 10% neutral formalin to fix overnight. The formalin was replaced with 70% ethanol prior to paraffin embedding. Tumors were counted under a stereotype microscope (Nikon Eclipse TE2000). The intestines were rolled up using a paper clip with a small loop to make Swiss rolls. The rolls were tied off on both ends using a 25-gauge syringe needle, cut in half, and placed in a paraffin cassette for embedding in paraffin for further analysis (H&E, immunohistochemical [IHC], or immunofluorescence [IF] staining).

The tissue sections were cut for hematoxylin (Agilent Dako) and eosin (Fisher Scientific) staining, and then H&E-stained sections were scanned using an Aperio digital pathology slide scanner (Leica Biosystems) for taking photomicrographs and counting tumors. Classification and grading of H&E-stained sections, including scoring tumor invasiveness, were performed by an experienced pathologist using an Olympus BX41 microscope.

### Primary organoid culture

Six-week-old PD mice and wild-type (WT) littermates (n = 6 mice per group) were killed, and their colons were harvested. Harvested colon tissues were digested with 10 mM ethylenediaminetetraacetic acid (EDTA) in chelation buffer (5.6 mM Na_2_HPO_4_, 8.0 mM KH_2_PO_4_, 96.2 mM NaCl, 1.6 mM KCl, 43.4 mM sucrose, 54.9 mM D-sorbitol, 0.5 mM DL-dithiothreitol) at room temperature for 10 min. Crypts were isolated by vigorously shaking the digested tissues for 30 s and then allowing sedimentation for 1 min prior to collecting the supernatant. The isolated crypts were counted using a hemocytometer and embedded in growth factor-reduced Matrigel (Cat# 356231, Corning) at 20 crypts/μL Matrigel. Approximately 20-μL droplets of Matrigel with crypts were seeded onto a flat-bottom 48-well plate (Cat# 3548, Corning). The Matrigel was solidified for 15 min in a 37°C incubator, and then 250 μL/well organoid culture medium (advanced Dulbecco’s modified Eagle medium/F12 supplemented with penicillin/streptomycin, 2 mM GlutaMAX, 10 mM HEPES [all from Invitrogen], 100 ng/mL mouse recombinant Wnt-3A [Millipore], 50 ng/mL mouse epidermal growth factor [Invitrogen], 100 ng/mL mouse recombinant noggin [Peprotech], 1 μg/mL human R-spondin-1 [Peprotech], 1 mM *N*-acetyl-L-cysteine [Sigma-Aldrich], 1× N-2 [Invitrogen], and 1× B-27 [Invitrogen]) was added. We added 10 μM Y-27632 (Sigma) to the media for the first 2 days’ culture, then changed the media without Y-27632 for later culture. The medium was replaced with fresh medium every 2 or 3 days. Organoids were maintained in an incubator with an atmosphere of 5% CO_2_. Organoids were counted, imaged, and quantified on day 7 of culture.

### IHC and IF staining

Paraffin-embedded tissue sections 5 μm thick were deparaffinized and rehydrated, and antigen retrieval was performed with an antigen unmasking solution (Vector Laboratory). The samples were then incubated in blocking buffer (PBS with 1.5% goat serum and 0.3% Triton X-100) for 1 h at room temperature and incubated with primary antibodies in a humidified chamber at 4°C overnight. For IHC staining, the following primary antibodies were used: β-catenin (Cat# ab32572, Abcam), Ki-67 (Cat# RM-9106-S1, Thermo Fisher), BMP7 (Cat# ab129156, Abcam), PPARD (Cat# ARP38765_T100, Aviva Systems Biology), phospho-rpS6(S235/236) (Cat# 4858S, Cell Signaling Technology), and phospho-TAK1(T184/T187) (Cat# 4508S, Cell Signaling Technology). The tissue sections were then incubated with biotinylated secondary antibodies (VECTASTAIN Elite ABC-HRP Kit, Cat# PK-6101, Vector Laboratories) for 1 h, followed by incubation with avidin-coupled peroxidase (Vector Laboratories) for 30 min. Diaminobenzidine (Agilent Dako) was used as the chromogen, and the slides were counterstained with Mayer’s hematoxylin (Agilent Dako).

To obtain a continuous score that took into account the IHC signal intensity and the frequency of positively stained cells in the human colon cancer tissue sections, we generated a combined expression score (CES) with a full range from 0 to 12. Staining intensity (SI) was scored as follows: 0 = no staining, 1 = light brown, 2 = brown, and 3 = dark brown. The percentage of positive cells (PP) was scored as follows: 0 = less than 10%, 1 = 10%-25%, 2 = 25%-50%, 3 = 50%-75%, and 4 = more than 75%. The CES for the nucleus or cytoplasm was calculated using the formula: CES = 4 (SI-1) + PP. The total PPARD CES was the CES of the combination of nucleus and cytoplasm.

For IF staining, the following primary antibodies were used: α-SMA (Cat# A5228, Millipore Sigma), E-cadherin (Cat# 710161, Thermo Fisher), and rRNA (Cat# sc-33678, Santa Cruz Biotechnology). The sections were washed 3 times in PBS-Tween 20 and incubated at room temperature for 2 h with Alexa Fluor 594-conjugated goat anti-rabbit immunoglobulin G (IgG) (H+L) secondary antibody (Cat# A-11012, Invitrogen) for E-cadherin or with Alexa Fluor 488-conjugated goat anti-mouse IgG (H+L) secondary antibody (Cat# A-11029, Invitrogen) for α-SMA and rRNA. The sections were washed 3 times in PBS-Tween 20 and mounted with ProLong Gold Antifade Mountant with 4′, 6-diamidino-2-phenylindole (Fisher Scientific).

### Generation of cell lines stably transfected with PPARD overexpression plasmid

The full length of the human PPARD cDNA was subcloned into a pcDNA3.1 plasmid (Life Technologies). HCT-116 cells were transfected with a pcDNA3.1 vector containing PPARD cDNA (PPARD vector) or pcDNA3.1 empty vector (control vector) and grown in selective medium containing hygromycin B (400 μg/mL, Roche Diagnostics). Stably transfected clones of PPARD vector-transfected (PPARD^SE^) and control vector-transfected cells were isolated and expanded.

### PPARD siRNA transfection

SW480 cells were cultured to 40% to 50% confluence and then transfected with 100 nM of 2 independent ON-TARGETplus PPARD small interfering RNAs (siRNAs) (Cat# J-003435-08-0002 and Cat# J-003435-09-0002, Dharmacon) or an equal amount of a nonspecific control siRNA (siGLO RISC-Free siRNA; Cat# D-001600-01-05, Dharmacon) using Lipofectamine 2000 (Invitrogen). The cells were harvested at 48 h after siRNA transfection for further analyses.

### Stable lentivirus transduction

Lentivirus plasmids for human PPARD (NM_006238) tagged ORF (Cat# RC214735L3, OriGene), mouse PPARD (NM011145) tagged ORF (Cat# MR207001L3, OriGene) in pLenti-C-myc-DDK-P2A-Puro, and a control lentivirus plasmid pLenti-C-Myc-DDK-P2A-Puro (Cat# PS100092, OriGene) were packaged into lentivirus particles (mouse PPARD, human PPARD, and control lentiviral particles) by MD Anderson’s shRNA and ORFeome Core Facility. Human (HCT-116 and SW480) or mouse (CT26) colorectal cancer cells were transduced with human PPARD or mouse PPARD or control lentiviral particles (10 MOIs for all lentiviruses) with hexadimethrine bromide (8 μg/mL). After 12 h, the culture medium was replaced with fresh medium containing puromycin (0.8 μg/mL for HCT-116, 4 μg/mL for SW480 and CT26). The medium was changed once every 48 to 72 h. Clones with stable transduction were isolated and expanded for further analyses.

### RNA extraction and quantitative real-time polymerase chain reaction

Mouse intestines were digested with 10 mM EDTA (pH 8.0) at room temperature for 10 min, and then digested crypts were harvested from the supernatants for further analysis. Total RNA was extracted from digested intestinal crypt cells from the mice or cancer cell lines using TRIzol for cell lines or an RNeasy Microarray Tissue Mini Kit (Qiagen) for tissue cells according to the manufacturer’s protocol. cDNA was synthesized using a Bio-Rad cDNA Synthesis Kit (Bio-Rad Laboratory). Quantitative real-time polymerase chain reaction (qRT-PCR) analyses were performed using a FastStart universal probe master (Roche) and StepOnePlus PCR system (Applied Biosystems). All the qRT-PCR probes were purchased from Applied Biosystems. See Table S2 for probe details. The relative RNA expression levels were normalized to the expression of mouse ACTB or human HPRT1 (Applied Biosystems), and calculated using a comparative threshold cycle method (ddC_t_).

### Western blotting analysis

Cells or digested intestinal crypts were homogenized in lysis buffer (0.5% Nonidet P-40, 20 mM 3-[N-morpholino] propanesulfonic acid [pH 7.0], 2 mM ethylene glycol tetraacetic acid, 5 mM EDTA, 30 mM sodium fluoride, 40 mM β-glycerophosphate, 2 mM sodium orthovanadate, 1 mM phenylmethylsulfonyl fluoride [Sigma], and 1× complete protease inhibitor cocktail [Roche Applied Science]). Fifty-microgram samples of each protein were separated onto 10-12% sodium dodecyl sulfate polyacrylamide gel. After electrophoresis, the proteins were transferred to a nitrocellulose membrane. The membranes were blocked with 5% milk or 5% bovine serum albumin for 2 h at room temperature and hybridized with primary antibodies at 4°C overnight. The following primary antibodies were used: BMP7 (for mouse/human, Cat# ab129156, Abcam), PPARD (for mouse, Cat# ab8937, Abcam), PPARD (for human, Cat# sc-7197, Santa Cruz Biotechnology), AKT1 (for human/mouse, Cat# 75692S, Cell Signaling Technology), phospho-AKT1(S473) (for human/mouse, Cat# 9018S, Cell Signaling Technology), AKT2 (for human/mouse, Cat# 3063S, Cell Signaling Technology), non-phospho/active β-catenin (S45) (for human/mouse, Cat# 19807S, Cell Signaling Technology), TAK1 (for human/mouse, Cat# 5206, Cell Signaling Technology), phospho-TAK1(T184/T187) (for human/mouse, Cat# 4508S, Cell Signaling Technology), PDGFRβ (for human/mouse, Cat# 3169S, Cell Signaling Technology), CDK1 (for human/mouse, Cat# 77055S, Cell Signaling Technology), EIF4G (for human/mouse, Cat# 2498S, Cell Signaling Technology), phospho-p38 MAPK(T180/Y182) (for human/mouse, Cat# 9211S, Cell Signaling Technology), phospho-rpS6(S235/236) (for human/mouse, Cat# 4858S, Cell Signaling Technology), phospho-GSK3β(S9) (for human/mouse, Cat# 5558, Cell Signaling Technology), β-actin (for human/mouse, Cat# sc-47778, Santa Cruz Biotechnology), and DDK (Cat# TA50011, OriGene). Next, the blots were hybridized with the secondary antibody for 1 h at room temperature. The blots were analyzed using enhanced chemiluminescence (GE Healthcare).

### Functional proteomics reverse phase protein array analysis

Reverse phase protein array (RPPA) analysis was performed on isolated intestinal epithelial cells from Apc^Δ580^ mice and their Apc^Δ580^-PD littermates according to the standard protocol at the RPPA Core Facility at MD Anderson. Specifically, relative protein levels for each sample were determined by interpolation of each dilution curve from the standard curve (supercurve) of the slide (antibody). The supercurve was constructed using a script in R written by the RPPA Core Facility. These values were defined as the supercurve log_2_ value. All data points were normalized for protein loading and transformed to a linear value, designated as “normalized linear.” The normalized linear value was transformed to a log_2_ value and then median-centered for further analysis. Median-centered values were centered by subtracting the median of all samples of a given protein. The information for the RPPA antibodies is available at https://www.mdanderson.org/research/research-resources/core-facilities/functional-proteomics-rppa-core/antibody-information-and-protocols.html. Student’s *t*-tests were used to assess differences in the expression levels of each gene between the Apc^Δ580^ and Apc^Δ580^-PD groups. In an exploratory analysis, volcano plots of the distributions of fold change (log_2_ [fold change]) for biological significance and Student’s *t*-test *P* values (-log_10_ [*P* value]) for statistical significance were used. We considered proteins as altered by PPARD overexpression if the absolute fold-change difference was greater than 2 or less than 0.5 and the Student’s *t*-test *P* value was less than 0.05; volcano plots were constructed.

### Chromatin immunoprecipitation-quantitative PCR assay

To determine whether PPARD binds to target gene (BMP7 or AKT1) promoters, we subjected SW480 cells stably transfected with human DDK-tagged PPARD or DDK-tagged control lentiviral particles to chromatin immunoprecipitation (ChIP). The cells were cross-linked by adding formaldehyde to the culture medium to a final concentration of 1% and incubating the medium for 10 min at 37°C. ChIP assays were performed using an EZ-ChIP Chromatin Immunoprecipitation Assay kit (Cat# 17-295, Millipore) according to the manufacturer’s protocol. Chromatin was immunoprecipitated using a specific anti-DDK antibody (Cat# TA50011, OriGene). We also used mouse IgG (Cat# sc-2762, Santa Cruz Biotechnology) as a negative control. The indicated gene promoters (-2000 to +500 bp, transcription start site was set as 0) were submitted to the online tool Genomatix MatInspector (https://www.genomatix.de/), and the calculated PPAR binding sites with matrix scores over 0.75 were considered significant. We then designed primers to amplify DNA fragments that contained each predicted PPARD binding site (pPDBS). The following primers were used: BMP7 (256 bp): 5′-CAGCAATTCCAGACAGCAAG-3′ (sense), 5′-CCCCTCAGTCCCTGTATCCT-3′ (antisense); AKT1 (127 bp): 5′-GTTGTCGGAGGAACTTCTGG-3′ (sense), reverse 5′-AAGTCCACTGGGAGGCAGAT-3′ (antisense). qRT-PCR was performed using SYBR Green ROX qPCR Mastermix (Cat# 330520, Qiagen) with amplification conditions as follows: 95°C for 10 min and then 95°C for 15 s and 60°C for 1 min, for 40 cycles. Input from each cell line was used as an endogenous control, and the relative enrichment level of IgG and DDK was calculated as 2^-ΔCt^. To assess the relative binding activity of PPARD to the BMP7 or AKT1 promoter region, the IgG enrichment value was normalized to 1, and relative enriched DDK values for each group were calculated based on the normalized IgG reading.

### Cell migration assay

For the migration assay, equal numbers of cells in culture medium with 0.5% FBS were plated on the tops of Corning Transwell inserts (8.0-μm pore), and 0.75 mL of chemical mixed whole medium was added to the lower compartment of the wells. After 36 h of incubation, cells that had not migrated were scraped from the top compartment, and cells that had migrated through the membrane (to the back of the membrane) were fixed and stained using the protocol of the HEMA 3 Stain Set (Thermo Fisher Scientific). Membranes were excised and mounted on a standard microscope slide (Curtin Matheson Scientific). The migrated cells in at least 4 random individual high-power fields per insert membrane were photographed with a light microscope and counted.

### β-catenin/TCF reporter assays

Cells were transfected with either pMegaTOP-FLASH reporter plasmid (14 WT TCF binding elements: CTTTGAT) or pMegaFOP-FLASH reporter plasmid (8 mutant TCF binding elements: CAAAGGG), and luciferase activity was measured as described previously.(50)

### Statistical analyses

Quantifiable outcome measures for 1 factor in experimental conditions were compared using 1-way analysis of variance, and Bonferroni adjustments were used for all multiple comparisons. We used 2-way analysis of variance to analyze data involving the simultaneous consideration of 2 factors. Tumor incidence was compared using chi-square tests. Survival rates as a function of time were estimated using the Kaplan-Meier method. The data were log-transformed as necessary to accommodate the normality and homoscedasticity assumptions implicit to the statistical procedures used. Data were analyzed using SAS software, version 9.4 (SAS Institute, Cary, NC) or GraphPad Prism 7.01. All tests were 2-sided and conducted at a significance level of *P* < 0.05.

## Acknowledgments

We thank Dr. Jail Park for his feedback on this work and Dr. Kenneth E. Hung at Tufts University for providing the Apc^Δ580^-flox mice. This work was supported by the National Cancer Institute (R01-CA142969, R01-CA195686, and R01-CA206539 to I.S.), the Cancer Prevention and Research Institute of Texas (RP140224 to I.S). We thank MS. Amy Ninetto at the Department of Scientific Publications at MD Anderson Cancer Center for editing the manuscript. This study made use of the MD Anderson Cancer Center Genetically Engineered Mouse Facility, Functional Proteomics Reverse Phase Protein Array Core, Sequencing and Microarray Facility, and Research Animal Support Facility at Houston, supported by Cancer Center Support Grant CA016672.

## Author contributions

I.S. developed the conceptual framework. I.S., X.Z., and Y.L. designed the experiments and wrote the manuscript. X.Z., Y.L, Y.D., R.T., L.W. W.C., W.X., F.L., S.G., J.C.J., S.P.C, and M.J.M performed experiments. X.Z., Y.L, Y.D., and I.S. analyzed the data. J.S.M. supervised and contributed to the statistical analyses. Y.D., M.G., and R.B. performed pathology analyses. D.W. provided conceptual feedback on the manuscript.

## Additional information

Supplemental Information includes 5 supplementary figures and 2 tables for experimental materials and can be found with this article online.

## REFERENCES

1. Morin PJ, Sparks AB, Korinek V, Barker N, Clevers H, Vogelstein B, and Kinzler KW. Activation of β-Catenin-Tcf Signaling in Colon Cancer by Mutations in β-Catenin or APC. Science. 1997;275(5307):1787–90.

2. Fearon ER, and Vogelstein B. A genetic model for colorectal tumorigenesis. Cell. 1990;61(5):759–67.

3. Oshima H, Nakayama M, Han TS, Naoi K, Ju X, Maeda Y, Robine S, Tsuchiya K, Sato T, Sato H, et al. Suppressing TGFbeta signaling in regenerating epithelia in an inflammatory microenvironment is sufficient to cause invasive intestinal cancer. Cancer Res. 2015;75(4):766–76.

4. Haggitt RC, Glotzbach RE, Soffer EE, and Wruble LD. Prognostic factors in colorectal carcinomas arising in adenomas: Implications for lesions removed by endoscopic polypectomy. Gastroenterology. 1985;89(2):328–36.

5. Michalik L, Desvergne B, and Wahli W. Peroxisome-proliferator-activated receptors and cancers: complex stories. Nat Rev Cancer. 2004;4(1):61–70.

6. He TC, Chan TA, Vogelstein B, and Kinzler KW. PPARdelta is an APC-regulated target of nonsteroidal anti-inflammatory drugs. Cell. 1999;99(3):335–45.

7. Gupta RA, Tan J, Krause WF, Geraci MW, Willson TM, Dey SK, and DuBois RN. Prostacyclin-mediated activation of peroxisome proliferator-activated receptor delta in colorectal cancer. Proc Natl Acad Sci U S A. 2000;97(24):13275–80.

8. Takayama O, Yamamoto H, Damdinsuren B, Sugita Y, Ngan CY, Xu X, Tsujino T, Takemasa I, Ikeda M, Sekimoto M, et al. Expression of PPAR[delta] in multistage carcinogenesis of the colorectum: implications of malignant cancer morphology. Br J Cancer. 2006;95(7):889–95.

9. Zuo X, Xu W, Xu M, Tian R, Moussalli MJ, Mao F, Zheng X, Wang J, Morris JS, Gagea M, et al. Metastasis regulation by PPARD expression in cancer cells. JCI Insight. 2017;2(1):e91419.

10. Foreman JE, Sorg JM, McGinnis KS, Rigas B, Williams JL, Clapper ML, Gonzalez FJ, and Peters JM. Regulation of peroxisome proliferator-activated receptor-β/δ by the APC/β-CATENIN pathway and nonsteroidal antiinflammatory drugs. Mol Carcinog. 2009;48(10):942–52.

11. Harman FS, Nicol CJ, Marin HE, Ward JM, Gonzalez FJ, and Peters JM. Peroxisome proliferator-activated receptor-delta attenuates colon carcinogenesis. Nat Med. 2004;10(5):481–3.

12. Wang D, Wang H, Guo Y, Ning W, Katkuri S, Wahli W, Desvergne B, Dey SK, and DuBois RN. Crosstalk between peroxisome proliferator-activated receptor delta and VEGF stimulates cancer progression. Proc Natl Acad Sci U S A. 2006;103(50):19069–74.

13. Han C, Lim K, Xu L, Li G, and Wu T. Regulation of Wnt/β-catenin pathway by cPLA2α and PPARδ. J Cell Biochem. 2008;105(2):534–45.

14. Scholtysek C, Katzenbeisser J, Fu H, Uderhardt S, Ipseiz N, Stoll C, Zaiss MM, Stock M, Donhauser L, Bohm C, et al. PPAR[beta]/[delta] governs Wnt signaling and bone turnover. Nat Med. 2013;19(5):608–13.

15. Beyaz S, Mana MD, Roper J, Kedrin D, Saadatpour A, Hong SJ, Bauer-Rowe KE, Xifaras ME, Akkad A, Arias E, et al. High-fat diet enhances stemness and tumorigenicity of intestinal progenitors. Nature. 2016;531(7592):53–8.

16. Xu M, Zuo X, and Shureiqi I. Targeting peroxisome proliferator-activated receptor-β/δ in colon cancer: How to aim? Biochem Pharmacol. 2013;85(5):607–11.

17. Peters JM, Gonzalez FJ, and Müller R. Establishing the Role of PPARβ/δ in Carcinogenesis. Trends Endocrinol Metab. 2015;26(11):595–607.

18. Hinoi T, Akyol A, Theisen BK, Ferguson DO, Greenson JK, Williams BO, Cho KR, and Fearon ER. Mouse Model of Colonic Adenoma-Carcinoma Progression Based on Somatic Apc Inactivation. Cancer Res. 2007;67(20):9721–30.

19. Schwitalla S, Fingerle Alexander A, Cammareri P, Nebelsiek T, Göktuna Serkan I, Ziegler Paul K, Canli O, Heijmans J, Huels David J, Moreaux G, et al. Intestinal Tumorigenesis Initiated by Dedifferentiation and Acquisition of Stem-Cell-like Properties. Cell. 2013;152(1):25–38.

20. Singh A, Sweeney Michael F, Yu M, Burger A, Greninger P, Benes C, Haber Daniel A, and Settleman J. TAK1 Inhibition Promotes Apoptosis in KRAS-Dependent Colon Cancers. Cell. 2012;148(4):639–50.

21. Shearer BG, Wiethe RW, Ashe A, Billin AN, Way JM, Stanley TB, Wagner CD, Xu RX, Leesnitzer LM, Merrihew RV, et al. Identification and characterization of 4-chloro-N-(2-{[5-trifluoromethyl)-2-pyridyl]sulfonyl}ethyl)benzamide (GSK3787), a selective and irreversible peroxisome proliferator-activated receptor delta (PPARdelta) antagonist. J Med Chem. 2010;53(4):1857–61.

22. Dolle RE, Dunn JA, Bobko M, Singh B, Kuster JE, Baizman E, Harris AL, Sawutz DG, Miller D, Wang S, et al. 5,7-Dimethoxy-3-(4-Pyridinyl)Quinoline Is a Potent and Selective Inhibitor of Human Vascular. beta.-Type Platelet-Derived Growth Factor Receptor Tyrosine Kinase. J Med Chem. 1994;37(17):2627–9.

23. Jackson RJ, Hellen CUT, and Pestova TV. The mechanism of eukaryotic translation initiation and principles of its regulation. Nature Reviews Molecular Cell Biology. 2010;11–113.

24. van de Wetering M, Sancho E, Verweij C, de Lau W, Oving I, Hurlstone A, van der Horn K, Batlle E, Coudreuse D, Haramis A-P, et al. The β-Catenin/TCF-4 Complex Imposes a Crypt Progenitor Phenotype on Colorectal Cancer Cells. Cell. 2002;111(2):241–50.

25. Yang X, Kume S, Tanaka Y, Isshiki K, Araki S-i, Chin-Kanasaki M, Sugimoto T, Koya D, Haneda M, Sugaya T, et al. GW501516, a PPARδ Agonist, Ameliorates Tubulointerstitial Inflammation in Proteinuric Kidney Disease via Inhibition of TAK1-NFκB Pathway in Mice. PLoS One. 2011;6(9):e25271.

26. Su X, Zhou G, Wang Y, Yang X, Li L, Yu R, and Li D. The PPARβ/δ Agonist GW501516 Attenuates Peritonitis in Peritoneal Fibrosis via Inhibition of TAK1–NFκB Pathway in Rats. Inflammation. 2014;37(3):729–37.

27. Stockert J, Wolf A, Kaddatz K, Schnitzer E, Finkernagel F, Meissner W, Müller-Brüsselbach S, Kracht M, and Müller R. Regulation of TAK1/TAB1-Mediated IL-1β Signaling by Cytoplasmic PPARβ/δ. PLoS One. 2013;8(4):e63011.

28. Grijelmo C, Rodrigue C, Svrcek M, Bruyneel E, Hendrix A, de Wever O, and Gespach C. Proinvasive activity of BMP-7 through SMAD4 /src -independent and ERK/ Rac /JNK -dependent signaling pathways in colon cancer cells. Cell Signal. 2007;19(8):1722–32.

29. Tan L, Gurbani D, Weisberg EL, Hunter JC, Li L, Jones DS, Ficarro SB, Mowafy S, Tam C-P, Rao S, et al. Structure-guided development of covalent TAK1 inhibitors. Bioorg Med Chem. 2017;25(3):838–46.

30. Gupta RA, Wang D, Katkuri S, Wang H, Dey SK, and DuBois RN. Activation of nuclear hormone receptor peroxisome proliferator-activated receptor-delta accelerates intestinal adenoma growth. Nat Med. 2004;10(3):245–7.

31. Grady WM. Genetic testing for high-risk colon cancer patients1 1Abbreviations used in this paper: FAP, familial adenomatous polyposis; HMPS, hereditary mixed polyposis syndrome; HNPCC, hereditary nonpolyposis colon cancer; JPS, juvenile polyposis; MMR, mutation mismatch repair; MSI, microsatellite instability; PJS, Peutz-Jeghers syndrome; TGF, transforming growth factor. Gastroenterology. 2003;124(6):1574–94.

32. Kitamura T, Kometani K, Hashida H, Matsunaga A, Miyoshi H, Hosogi H, Aoki M, Oshima M, Hattori M, Takabayashi A, et al. SMAD4-deficient intestinal tumors recruit CCR1+ myeloid cells that promote invasion. Nat Genet. 2007;39(4):467–75.

33. Brabletz T, Jung A, Reu S, Porzner M, Hlubek F, Kunz-Schughart LA, Knuechel R, and Kirchner T. Variable beta-catenin expression in colorectal cancers indicates tumor progression driven by the tumor environment. Proc Natl Acad Sci U S A. 2001;98(18):10356–61.

34. Wu Z-Q, Brabletz T, Fearon E, Willis AL, Hu CY, Li X-Y, and Weiss SJ. Canonical Wnt suppressor, Axin2, promotes colon carcinoma oncogenic activity. Proceedings of the National Academy of Sciences. 2012;109(28):11312–7.

35. Sillars-Hardebol A, Carvalho B, de Wit M, Postma C, Delis-van Diemen P, Mongera S, Ylstra B, van de Wiel M, Meijer G, and Fijneman RA. Identification of key genes for carcinogenic pathways associated with colorectal adenoma-to-carcinoma progression. Tumor Biol. 2010;31(2):89–96.

36. Hay N, and Sonenberg N. Upstream and downstream of mTOR. Genes Dev. 2004;18(16):1926–45.

37. Di-Poi N, Ng CY, Tan NS, Yang Z, Hemmings BA, Desvergne B, Michalik L, and Wahli W. Epithelium-Mesenchyme Interactions Control the Activity of Peroxisome Proliferator-Activated Receptor {beta}/{delta} during Hair Follicle Development. Mol Cell Biol. 2005;25(5):1696–712.

38. Pedchenko TV, Gonzalez AL, Wang D, DuBois RN, and Massion PP. Peroxisome proliferator-activated receptor beta/delta expression and activation in lung cancer. Am J Respir Cell Mol Biol. 2008;39(6):689–96.

39. Yuan H, Lu J, Xiao J, Upadhyay G, Umans R, Kallakury B, Yin Y, Fant ME, Kopelovich L, and Glazer RI. PPARdelta induces estrogen receptor-positive mammary neoplasia through an inflammatory and metabolic phenotype linked to mTOR activation. Cancer Res. 2013;73(14):4349–61.

40. McCubrey JA, Steelman LS, Bertrand FE, Davis NM, Sokolosky M, Abrams SL, Montalto G,D’Assoro AB, Libra M, Nicoletti F, et al. GSK-3 as potential target for therapeutic intervention in cancer. Oncotarget. 2014;5(10):2881–911.

41. Kapoor A, Collino M, Castiglia S, Fantozzi R, and Thiemermann C. ACTIVATION OF PEROXISOME PROLIFERATOR-ACTIVATED RECEPTOR-β/δ ATTENUATES MYOCARDIAL ISCHEMIA/REPERFUSION INJURY IN THE RAT. Shock. 2010;34(2):117–24.

42. Kapoor A, Shintani Y, Collino M, Osuchowski MF, Busch D, Patel NSA, Sepodes B, Castiglia S, Fantozzi R, Bishop-Bailey D, et al. Protective Role of Peroxisome Proliferator–activated Receptor-β/δ in Septic Shock. Am J Respir Crit Care Med. 2010;182(12):1506–15.

43. Ruggero D. Translational control in cancer etiology. Cold Spring Harb Perspect Biol. 2013;5(2).

44. Boussemart L, Malka-Mahieu H, Girault I, Allard D, Hemmingsson O, Tomasic G, Thomas M, Basmadjian C, Ribeiro N, Thuaud F, et al. eIF4F is a nexus of resistance to anti-BRAF and anti-MEK cancer therapies. Nature. 2014;513–105.

45. Enserink JM, and Kolodner RD. An overview of Cdk1-controlled targets and processes. Cell Div. 2010;5(1):11.

46. Shuda M, Velásquez C, Cheng E, Cordek DG, Kwun HJ, Chang Y, and Moore PS. CDK1 substitutes for mTOR kinase to activate mitotic cap-dependent protein translation. Proceedings of the National Academy of Sciences. 2015;112(19):5875–82.

47. Brown NR, Korolchuk S, Martin MP, Stanley WA, Moukhametzianov R, Noble MEM, and Endicott JA. CDK1 structures reveal conserved and unique features of the essential cell cycle CDK. Nature Communications. 2015;6(6769.

48. Zuo X, Xu M, Yu J, Wu Y, Moussalli MJ, Manyam GC, Lee SI, Liang S, Gagea M, Morris JS, et al. Potentiation of colon cancer susceptibility in mice by colonic epithelial PPAR-delta/beta overexpression. J Natl Cancer Inst. 2014;106(4):dju052.

49. Hung KE, Maricevich MA, Richard LG, Chen WY, Richardson MP, Kunin A, Bronson RT, Mahmood U, and Kucherlapati R. Development of a mouse model for sporadic and metastatic colon tumors and its use in assessing drug treatment. Proceedings of the National Academy of Sciences. 2010;107(4):1565–70.

50. Park J-I, Venteicher AS, Hong JY, Choi J, Jun S, Shkreli M, Chang W, Meng Z, Cheung P, Ji H, et al. Telomerase modulates Wnt signalling by association with target gene chromatin. Nature. 2009;460(7251):66–72.

